# LipidRT: leveraging retention time coherence for robust lipid annotation in reversed-phase LC-MS-based lipidomics

**DOI:** 10.64898/2026.11.11.765883

**Authors:** Celia Regal-Marqués, María Isabel Alcoriza-Balaguer, Hana Cermakova, Marta Roca, Óscar Juan-Vidal, Julián Carretero, Ana Gradillas, Coral Barbas, Juan Carlos García-Cañaveras, Agustín Lahoz

## Abstract

MS/MS data alone is often insufficient for confident lipid identification, while existing retention time (RT)-based methods, despite their promise, have limited applicability. We present LipidRT, an R-based Shiny application that uses RT coherence to validate, refine and expand lipid annotations in untargeted RPLC-HRMS lipidomics in a dataset-specific, software-agnostic manner. By exploiting conserved chromatographic elution relationships as structural constraints and excluding RT-inconsistent annotations from model fitting, LipidRT improves annotation confidence, coverage and consistency while minimizing manual curation.

---

Liquid chromatography (LC) coupled with high-resolution mass spectrometry (HRMS) is the leading analytical approach for untargeted lipidomics. Lipid annotation relies on MS/MS spectra and spectral matching or fragmentation rules to infer lipid structures, ranging from sum composition [e.g. PC(34:1)] to full structural elucidation, including double-bonds (DB) location and *cis*/*trans* geometry [e.g. PC(16:0/18:1(9*Z*))]^1^. Automated MS/MS-based annotation remains challenging because: i) isobaric and isomeric lipids from different subclasses often generate similar fragments, leading to ambiguous identifications; ii) adduct formation and in-source fragmentation vary with experimental conditions, analytical platforms and biological matrices; iii) co-eluting lipids produce convoluted spectra; and iv) low-abundance lipids provide limited structural information^2^.

In reverse-phase LC (RPLC), lipid elution follows predictable patterns^3^. Retention time (RT) modeling based on equivalent carbon number (ECN)^3,4^ and Quantitative Structure-Retention Relationships (QSRR) combined with machine learning^5,6^ have improved lipid identification confidence and have been implemented in tools such as LDA^7^or MS-DIAL^8^, and more recently into LipidIN^9^. However, important limitations remain: i) RT models are often developed for specific chromatographic conditions, reducing transferability across laboratories and platforms; ii) models fitted directly to annotated datasets can be compromised by incorrect initial identifications which may distort RT relationships (Supplementary Fig. 1, Supplementary Note 1, Supplementary Table 1); iii) some isobaric/isomeric species cannot be reliably distinguished by MS/MS alone, requiring additional rules (e.g. PC(O-18:1/20:4) and (PC(P-18:0/20:4)). Therefore, careful manual review integrating MS/MS spectra, adduct profiles, RT and expected fatty acyl/alkyl/alkenyl chain composition remains the most reliable strategy for comprehensive lipidome characterization^2,10^.

We hypothesize that the structural and chromatographic relationships governing lipid retention in RPLC are sufficiently conserved to be applied as an independent layer, downstream of any annotation software, to improve annotation accuracy and coverage across RPLC platforms.

Here, we introduce LipidRT, an R-based Shiny application that exploits RT coherence to validate and extend lipid annotation in untargeted RPLC-MS datasets. LipidRT first applies structure-based elution rules to remove annotations with RTs inconsistent with their expected chromatographic behavior. The resulting high-confidence set is used to build dataset-specific RT models that score existing annotations and recover previously unidentified lipid species based on MS1 information (*m/z* and RT) (Fig. 1a). Because it requires only a feature table containing *m/z*, RT, lipid name and adduct information, LipidRT can be applied to the output of any annotation software. A detailed description of the adducts (Supplementary Table 2), lipid classes/subclasses (Supplementary Table 3) and fatty acyl chains (Supplementary Table 4) supported by LipidRT, and how to expand them is provided in Supplementary Note 2. The workflow can run fully automatically, while the Shiny interface enables interactive review and curation of results.

**Fig. 1.**
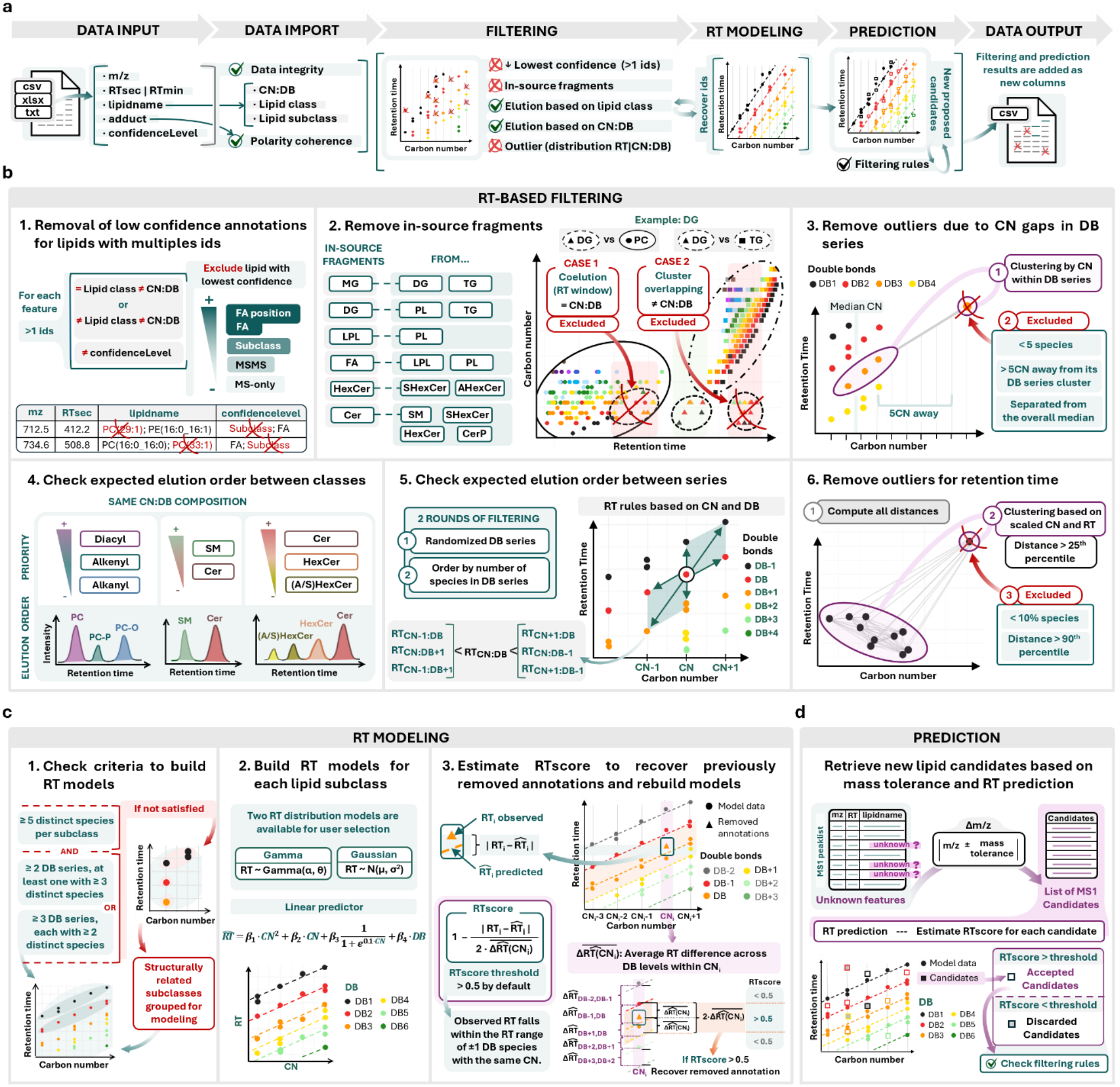
LipidRT workflow. **a**, Overview of the LipidRT workflow, from data import and input requirements to results export. **b**, Six-step filtering algorithm used to remove RT-inconsistent annotations before model fitting. **c**, RT-based modeling and scoring. **d**, Prediction of novel lipid annotations.

RT models must be built from high-confidence annotations to accurately capture chromatographic elution patternsas shown in Supplementary Note 1 using the in-house dataset QExactive_DS1 (generated using the NIST® Standard Reference Material® 1950 human plasma (SRM 1950) acquired on an Orbitrap Q Exactive^−^Plus mass spectrometer). We therefore implemented a six-step filtering algorithm to remove RT-inconsistent annotations before model fitting (Fig. 1b). First, for features with multiple annotations, only the one with the highest structural resolution is retained. Second, in-source fragments (e.g., DG from co-eluting TG or PC) are identified and removed. Third, species separated by large carbon number (CN) gaps within a DB series are discarded. Fourth, the elution order of related classes (e.g., SM and Cer) is evaluated and suspicious annotations are removed. This step is particularly important for diacylglycerophospholipids (PL), plasmenyl (PL(P-)) and plasmanyl (PL(O-)) lipids, which are often indistinguishable by MS/MS alone^11,12^. RT therefore provides a critical orthogonal criterion, following the relationship: RT[PL(O-CN:DB+1)] < RT[PL(CN:DB)] < RT[PL(P-CN:DB)] < RT[PL(O-CN:DB)] (e.g., PC(O-18:1/20:4) < PC (18:0/20:4) < PC(P-18:0/20:4) < PC(O-18:0/20:4) (Supplementary Note 3, Supplementary Fig. 2, Supplementary Tables 5-6). Fifth, each lipid is compared with neighboring species in CN:DB space and annotations violating the expected elution order are removed. Finally, outliers in the C-RT space within each class are excluded. Further details are provided in Methods.

For the retained annotations, RT models are fitted for each lipid class using a generalized linear model (Gamma by default) with quadratic and sigmoidal CN terms and a linear DB term, constrained to preserve the expected elution order (Fig. 1c). When insufficient annotations are available, structurally related classes are combined and lipid class is included as a predictor. The fitted models estimate theoretical RT values, from which an RTscore is calculated by scaling the deviation between observed and predicted RT to the spacing between adjacent DB series. Previously discarded annotations are re-evaluated and reinstated when their RTscore exceeds 0.5. Then, unannotated MS1 features are matched to candidate lipids based on *m/z* and expected adducts, and candidates with RTscores > 0.5 are proposed as new annotations, ranked by RTscore when multiple assignments are possible. Finally, proposed annotations are verified against key RT-based rules (Fig. 1d). Further details are provided in Methods.

We first evaluated the automated LipidRT workflow using the manually curated LC-MS lipidomics database of SRM 1950 reported Martinez *et al*., 2024^10^ as reference (Agilent_DS2). To assess filtering performance, we introduced artificial annotations with progressively larger RT deviations across lipid subclasses. LipidRT increasingly removed these annotations as chromatographic inconsistency increased, from ∼70% to ∼87% across deviations ranging from ±1.5 to ±4.5 expected RT variation, largely independent of the proportion introduced (Fig. 2a, Supplementary Table 7). To assess prediction performance, 5-40% of high-confidence annotations were randomly removed within each lipid subclass. LipidRT recovered ∼89% of omitted annotations, while ∼98% of predictions were correct, indicating a conservative and highly accurate prediction strategy (Fig. 2a, Supplementary Table 8).

**Fig. 2.**
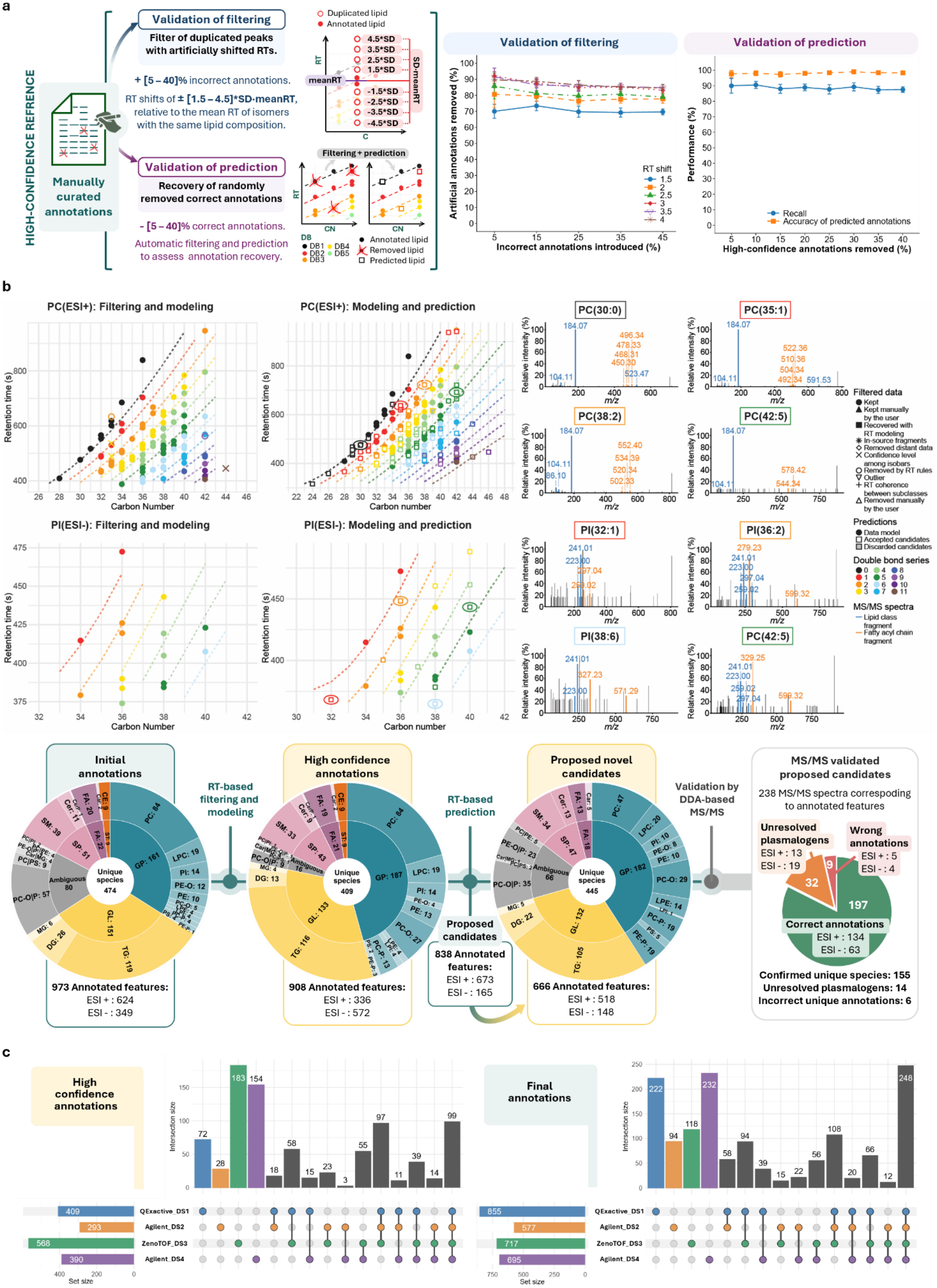
Validation of LipidRT using NIST SRM 1950 human plasma. **a**, In silico validation of the LipidRT filtering and prediction algorithms by introducing erroneous annotations or removing high-confidence annotations from the manually curated database reported by *Martinez et al 2024* ^10^ (Agilent_DS2). **b**, Application of the LipidRT workflow to the Qexactive_DS1 dataset, including MS/MS validation of newly proposed annotations. **c**. Comparison of lipidome coverage across four RPLC-MS datasets before and after LipidRT-assisted curation. QExactive_DS1 corresponds to an in-house generated dataset on an Orbitrap Q Exactive^−^ Plus; Agilent_DS2 was described in *Martinez et al. 2024*^10^; ZenoTOF_DS3, in *Martinez et al. 2025*^14^; and Agilent_DS4, in *Tsugawa et al. 2020*^8^.

As an additional experimental validation of LipidRT, we applied the workflow to the QExactive_DS1 annotated with LipidMS^13^ (Fig. 2b and Supplementary Note 5). Following the complete workflow, including manual inspection, LipidRT reduced suspicious and ambiguous annotations while substantially expanding lipid coverage. The number of unique lipid species more than doubled, including an 80% increase in unique CN:DB combinations, largely by completing gaps within lipid series (Supplementary Table 9). Iterative DDA acquisitions validated the proposed identities: of 238 candidates yielding MS/MS spectra, 197 (83%) confirmed the proposed annotation (193 as the top-ranked candidate), 32 supported the ether lipid assignment but not the O/P-isomerism, and only 9 (4%) were inconsistent with the prediction (Supplementary Table 10).

We next evaluated the ability of LipidRT to support expert curation and harmonize lipid annotations across inhouse and published^8,10,14^ SRM 1950 datasets generated on different RPLC-HRMS platforms and processed with distinct annotation workflows (Supplementary Tables 11-12). Following automated LipidRT analysis and manual review, ∼18% of initial annotations were removed, whereas only ∼6% of retained features required further curation, substantially reducing expert workload (Supplementary Table 13). Although the initial lipidome profiles showed limited agreement, with only 99 unique species shared across all four datasets, LipidRT-assisted curation and coverage expansion increased this overlap to 248 species (150% increase), while increasing the total number of annotated species by 58% (Fig. 2c, Supplementary Note 6 and Supplementary Fig. 6).

LipidRT provides a robust RT-based framework for improving the accuracy, consistency and coverage of lipid annotations in untargeted RPLC-MS lipidomics. By combining RT-based filtering with dataset-specific modeling, it increases confidence in both existing and newly proposed annotations. Validation across SRM 1950 datasets demonstrated high recovery and prediction accuracy, expanded lipid coverage, strong agreement with MS/MS confirmation, and improved annotation harmonization across studies. Together with its interactive Shiny interface, LipidRT reduces expert curation effort while enabling more comprehensive and reproducible lipidome characterization.

## Methods

### Development of LipidRT

#### Implementation

LipidRT (v0.1.0) has been built as a R package using R version (4.5.1)^15^ and it is available at the Comprehensive R Archive Network (CRAN). It has been designed as a Shiny application to facilitate interactive data inspection and manual curation, providing an intuitive interface for the lipidomics community^16^.

#### Data input requirements

The input for LipidRT consists of a peak table of lipid annotations and unannotated features provided in a tabular format. This table can be supplied as a .txt, .csv, or .xlsx file, or as a data frame directly in R. The peak table may originate from different lipid annotation software tools. For inputs that do not originate from LipidMS or MS-DIAL, the table must strictly include the following columns with these exact names: “*mz*”, “*RTsec*” or “*RTmin*”, “*lipidname*” and “*adduct*” and the column “*confidenceLevel*” is recommended although it can be inferred based on the information provided in “*lipidname*”. The column “*mz*” represents the mass-to-charge ratio of the detected feature, while “*lipidname*” contains the lipid annotation according to the common name nomenclature (e.g., PC(16:0/18:1), PC(34:1), for features with more than one annotation, they have to be separated by the symbol “|”). The “*adduct*” column specifies the ion adduct type. Retention time (RT) must be provided as either “*RTsec*” or “*RTmin*”, depending on whether the values are expressed in seconds or minutes. Finally, the column “*confidenceLevel*” is recommended to indicate the reliability of the annotation. Accepted confidence levels follow a hierarchical order from lowest to highest: MS-only, MSMS, Subclass, FA, and FA position. If this column is missing, the workflow will attempt to infer the confidence level from the lipid annotations during processing. These column names are mandatory for proper function execution, and any deviation will prevent the data from being correctly imported. Optionally, a column named “*IS*”, to indicate whether a lipid is an internal standard, can be included. LipidRT only supports one polarity at a time and thus datasets containing simultaneous annotations from both positive and negative ionization modes, inferred by the information contained in the “*adduct*” column, cannot be processed.

#### Data import

Upon importing a peak table, LipidRT first attempts to identify its source, which can be LipidMS, MS-DIAL, or other, based on software-specific column names within the table header. After validating the table structure, LipidRT ensures that all relevant variables have the appropriate data type, converting numerical fields when necessary, and subsequently standardizes the peak table format across the supported software platforms. This harmonization step generates a consistent data structure that can be used in all downstream filtering and retention time prediction analyses.

The standardized table is initially built from the required columns described above. In addition, a new column named *LipidRTid* is created to uniquely identify each feature. The software then checks whether a *confidenceLevel* column is available. For LipidMS and *Other* peak tables, an existing *confidenceLevel* column is retained without modification. In contrast, for MS-DIAL peak tables, the confidence level is derived from the *Annotation tag (VS1*.*0)* column and translated into the LipidRT nomenclature. Specifically, the values *“Lipid assigned”* and *“Chain resolved”* are converted to *“FA position”, “Chain semi-resolved”* is converted to *“FA”, “Class resolved”* is converted to *“Subclass”*, and *“Annotated by experimental MS/MS”, “annotated by user-defined text library”*, and *“Unknown”* are converted to *“MSMS”*. If a *confidenceLevel* column is not available, LipidRT infers the confidence level directly from the lipid annotation. Annotations containing fatty acyl chains separated by “/” are assigned to the confidence level *“FA position”*, annotations containing chains separated by “_” are assigned *“FA”*, and annotations reported only as total composition, without resolved fatty acyl chains, are assigned *“Subclass”*.

Following standardization, LipidRT parses the lipid annotations associated with each feature. When multiple lipid annotations are reported for the same feature, they are expected to be separated by the “|” character. Each annotation is extracted and stored as an independent entry, while retaining the same *LipidRTid* to indicate that they originate from the same feature. A new column named *index* is generated to uniquely identify each annotation. During this process, key information is extracted from each annotation, including the lipid class, lipid subclass, total number of carbon atoms, number of double bonds, and adduct. These variables are subsequently used during filtering and retention time prediction.

An additional column named *Filter* is then generated to track the filtering status of each annotation. Annotations labelled as *“Kept”* are those that will be considered during the subsequent filtering steps. In some cases, a feature may contain multiple annotations corresponding to the same lipid species but associated with different confidence levels. To avoid overrepresentation of such annotations during filtering, only the annotation with the highest confidence level is labelled as *“Kept”*, while the remaining duplicated annotations are assigned the label *“Ignore duplicated”*. These duplicated annotations are temporarily excluded from the filtering procedure but are restored after filtering is completed, inheriting the filtering outcome assigned to the retained annotation.

The *Filter* column may also contain the value *“Not Pass QC”*, which is assigned to annotations beginning with *“No MS2”* or *“low score”*. These annotations are considered low-confidence identifications and are therefore excluded from filtering. However, during the retention time prediction stage, the corresponding features are treated as candidates for reannotation, allowing LipidRT to propose alternative lipid candidates based on their *m/z* values. Finally, annotations assigned the value *“Unsupported class”* correspond to lipid classes that cannot currently be recognized by LipidRT (Supplementary Table 3). These annotations are not subjected to filtering but are retained until the end of the analysis. Furthermore, features associated with unsupported classes are not considered candidates for retention time prediction.

Once the annotations have been standardized and expanded into a data frame where each row corresponds to a single annotation, with the lipid class, subclass, CN, DB, and adduct explicitly extracted, the polarity of the dataset is determined based on the detected adducts. If only positive-mode adducts are present, the analysis is assigned positive polarity; similarly, if only negative-mode adducts are detected, the analysis is assigned negative polarity. When adducts from both ionization modes are present, the dataset is classified as having mixed polarity. In such cases, downstream filtering and retention time prediction analyses are not performed, as LipidRT requires data acquired in a single ionization polarity.

#### RT-based filtering

The main six-step algorithm of the filtering workflow implemented in LipidRT include:

1. Removal of low confidence annotations for lipids with multiples ids. For features with proposed annotations from different lipid classes and/or adducts, only the assignment with the highest annotation confidence (i.e. higher structural characterization) is retained. For example, features simultaneously annotated as MG and carnitines are reviewed, with the latter being retained since carnitines generally exhibit clearer MS/MS fragmentation patterns. Ether lipids (PL(O-)/PL(P-)) are also evaluated to prevent misannotations between phospholipids (PL) and lysophospholipids (LPL); in these cases, LPL annotations are prioritized when fragments of fatty acyl chains supporting the PL annotation are missing. In addition, for ether lipids, when several annotations are proposed for a single feature, only the one with the highest confidence level is retained.
2. Removal of in-source fragments. In-source fragments are detected using RT-based information and by evaluating overlaps between lipid classes and their corresponding fragment classes, such as MG from DG and TG, DG from TG or PL, and LPL from PL. When a potential in-source fragment shares the same CN and DB composition as the parent lipid [e.g. DG(34:1) and PC(34:1)], coelution is verified within a user-defined RT window. For fragments with different carbon and double-bond compositions [e.g. DG(34:1) and TG(52:2)], candidate species are clustered by RT and CN using a stepwise merging algorithm. Both variables are scaled and weighted, with greater emphasis on RT. Clusters are iteratively merged until their distance exceeds a threshold (default: 25^th^ percentile of all pairwise distances calculated with a single linkage algorithm). Overlap between fragment and reference lipid classes is then evaluated, and clusters showing more than 75% overlap with 95^th^ percentile of the reference class RT distribution are marked as in-source fragments.
3. Removal of outliers due to CN gaps in DB series. Within each lipid class, species that show a substantial deviation in CN from the main cluster of their corresponding DB series are identified by computing the pairwise distance in CN composition and applying a user-defined threshold. Clusters are defined within each DB series using a CN gap specified by the user (5 carbons by default) [e.g. PC(26:0) would be removed if the next saturated PC in the series is PC(32:0)]. To prevent excessive filtering, a maximum number of species that can be removed is also defined. By default, the outlier criteria correspond to a carbon-distance threshold of 5 and a maximum removable cluster size of five species.
4. Check expected elution order between related classes. First, ether phospholipids are evaluated using their corresponding diacyl species as reference. When the “*include all isobaric species”* option is enabled, all compatible isomeric O- and P-species are considered. Species with resolved fatty acyl chains are compared only with their homologous diacyl species, whereas annotations at total composition level are compared with all diacyl species sharing the same composition. The three RT rules applied to ether phospholipids are detailed in Supplementary Note 3. Second, sphingolipids are evaluated using dihydroxy-base species (SL_d) as reference. The expected relationships for the RT are: (i)SL_t < SL_d; (ii) SL(t_X) < SL(d_X+1); and (iii) SHexCer < HexCer < Cer < AHexCer. Reference SL_d species are retained when these rules are not satisfied. Finally, DGDG, MGDG and DG are evaluated according to the expected order DGDG < MGDG < DG, with DG used as the reference and retained in case of inconsistency.
5. Check expected elution order between CN:DB series for a lipid class. For each lipid species, the following conditions must be satisfied based on the median of the RT of all the isobaric/isomeric species being compared: (i) lipids with fewer carbons and the same DB number must elute earlier; (ii) lipids with the same CN and more double bonds must elute earlier; (iii) lipids with more carbons and the same DB number must elute later; (iv) lipids with the same CN and fewer double bonds must elute later; (v) lipids with more carbons and fewer double bonds must elute later; and (vi) lipids with fewer carbons and more double bonds must elute earlier. Because the order in which species are evaluated may influence the outcome, these rules are applied twice. In the first round, species are reviewed in randomized DB series order and removed if they are flagged in at least 80% of iterations. In the second round, DB series containing more species are evaluated first, thereby prioritizing the most populated series.
6. Removal of outliers based on retention time clustering. Outliers in RT are detected by clustering species within each lipid class, using a maximum distance corresponding to the 25th percentile of all pairwise RT-based distances. Clusters representing less than 10% of the total class and separated by more than the 90th percentile of distances are considered RT outliers and removed.

#### RT modeling

RT for each lipid class is fitted using a second-order polynomial predictor that includes the number of carbons (CN^2^ and CN), the number of double bonds (DB), and a smoothing term to stabilize the fit at the late-eluting region of the chromatogram, which can introduce local curvature. At the lower end of the carbon range, this polynomial predictor may introduce an artificial decrease in RT with increasing CN, while at the upper end, predicted times may decrease after a local maximum. Both behaviors do not follow the theoretically expected elution pattern. To correct this, RT for lipids below a local minimum are adjusted to follow a monotonic increasing slope based on the difference between the minimum and the preceding lipid, while for lipids above a local maximum, retention times are capped at the value of the maximum. These adjustments ensure that predicted RT increase consistently with CN and degree of unsaturation while preserving the overall structure of the predictive model. The general form of the model is:

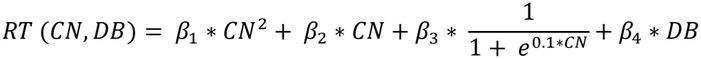

Models were fitted using the glm() function in R, with either a Gaussian family and identity link or a Gamma family with log link, as specified by the user. By default, LipidRT uses the Gamma distribution, and the fitted model is:

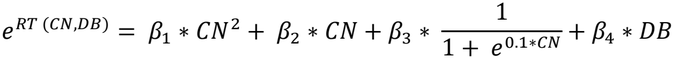

Lipid classes are modeled only when sufficient data are available: a) at least five unique CN:DB combinations distributed across two DB series, with at least one series containing three or more unique CN:DB combinations; b) at least six unique CN:DB combinations distributed in at least three DB series with at least two unique CN:DB combinations each. If these criteria are not met, a combined model is constructed using data from structurally related lipid subclasses, incorporating the lipid subclass factor itself as an additional predictor variable:

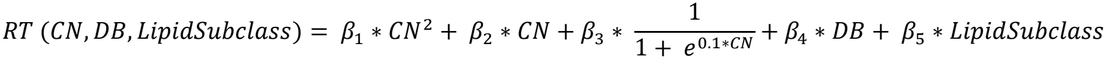

For the Gamma distribution the modified equation is:

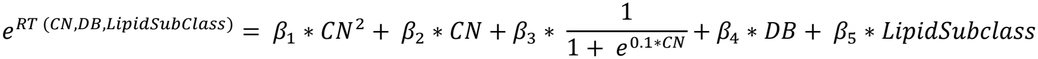

Specifically, all diacyl phospholipids are grouped together (PC, PE, PS, PI, PG and PA); their corresponding lysophospholipids too (LPC, LPE, LPS, LPI, LPG and LPA), ether-linked subclasses are also grouped by headgroup (PC, PC-O and PC-P on one side and PE, PE-O and PE-P on the other); SM_d/t are grouped with Cer_d/t if required; ceramides are also grouped together (Cer_d, Cer_t, HexCer_d, HexCer_t, SHexCer_d, SHexCer_t, AHexCer_d and AHexCer_t); and DG, MGMG, DGDG are also grouped together.

To evaluate the agreement between the observed and predicted retention times, an RT prediction score (*RTscore*) is calculated for each lipid species. The score quantifies how well the experimentally measured RT matches the value predicted by the class-specific RT model, normalized by the expected retention-time variability for species with the same CN:DB composition. The *RTscore* for lipid *i* is defined as:

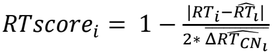

where *RT*_*i*_ is the observed retention time, 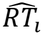 is the predicted retention time, and 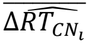 corresponds to the expected RT variability for lipids with the same CN as lipid *i*, calculated as the media of all differences between predicted and observed RT for those lipid species. 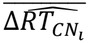 provides a class- and composition-specific normalization factor, preventing systematic biases between lipid subclasses. The raw score can theoretically range from −∞ to 1, however, negative values are truncated to 0 to ensure interpretability. Therefore, RTscores range from 0 to 1, with values closer to 1 indicating higher agreement between observed and predicted RT. Lipid annotations with RTscore > 0.5 are considered consistent with the model and retained as a value of 0.5 is expected for the RT of a lipid belonging to the series differing 1DB with respect to the series being evaluated. Previously discarded annotations are re-evaluated and reinstated if their RTscore exceeds 0.5 or the user-defined threshold.

#### Prediction of novel annotations

All potential missing annotations are identified from MS1 features according to their *m/z* values and expected adducts for each lipid class. Theoretical RT values are estimated for all candidate features, and those with RTscores > 0.5 are proposed as new annotations. To prevent false positive annotations, the following RT-based rules are checked and only those new annotations that adhere to them are finally retained as accepted candidates: i) gap in CN:DB series; ii) elution order between classes; and iii) elution order within classes.

#### Data export

Once an analysis is completed, all results are stored in the *rtdata* object, including filtering results and, when performed, RT models and predicted annotations.

Filtered, predicted and final lipid annotations are added to the original feature table provided as input, while retaining all original features and associated information. LipidRT also generates a summarized feature table containing the original feature information together with the final lipid annotations resulting from the complete workflow.

The complete *rtdata* object can be exported for further analysis in R. In addition, output tables, filtering and prediction plots, RT model reports, and predicted lipid species plots can be downloaded directly from the LipidRT Shiny application.

### Materials

Isopropanol (IPA), ammonium formate, and ammonium acetate used for sample preparation and LC-MS analysis were purchased from Sigma-Aldrich/Fluka (Madrid, Spain). Acetonitrile (ACN) was obtained from Fisher Scientific (Loughborough, UK). The deuterated lipid standard mix (SPLASH^®^ LIPIDOMIX^®^) was purchased from Avanti Polar Lipids (Alabaster, AL, USA). The human plasma NIST® Standard Reference Material® 1950 (SRM 1950) was purchased from Merck (Madrid, Spain).

### Sample preparation

For lipid extraction, 50 µL of the SRM 1950 were mixed with 150 µL of IPA containing the deuterated internal standards at 0.5 µg/mL each. After vortexing, samples were incubated for 20 min at −20 °C and then centrifuged for 15 min at 15,000 *g* and 4 °C. Finally, 100 µL of the supernatant were transferred into HPLC vials for LC–MS analysis^17^.

### RP-UHPLC-MS analysis

Lipidomic profiling was performed on a Quadrupole–Orbitrap mass spectrometer (Q Exactive^−^ Plus, Thermo Fisher Scientific) coupled to reverse-phase liquid chromatography (RP-LC) via electrospray ionization (ESI)^13^. Chromatographic separation was achieved using a CSH C18 column (2.1 × 100 mm, 1.7 µm particle size; Waters). Solvent A consisted of 10 mM ammonium acetate (for ESI(+)) or ammonium formate (for ESI(–)) in ACN/H_2_O (60:40, v/v), whereas solvent B contained 10 mM ammonium acetate (for ESI(+)) or ammonium formate (for ESI(–)) in IPA/ACN (90:10, v/v). The flow rate was 0.4 mL/min, the column temperature was maintained at 65 °C, the autosampler at 5 °C, and the injection volume was 5 µL. The LC gradient was^13^: 0 min, 20% B; 2 min, 20% B; 4 min, 43% B; 4.1 min, 50% B; 14 min, 54% B; 14.1 min, 70% B; 20 min, 99% B; 24 min, 99% B; 24.5 min, 20% B; and 27 min, 20% B. The following source conditions were applied for ESI(+): sheath gas flow rate, 25 (0–14 min) and 80 (14– 27 min); auxiliary gas flow rate, 10 (0–14 min) and 25 (14–27 min); spray voltage, 3.0 kV; capillary temperature, 215 °C; ^−^); S-lens RF level, 95; and auxiliary gas heater temperature, 215 °C. For ESI(–) the source conditions were: sheath gas flow rate, 25; auxiliary gas flow rate, 25; spray voltage, 2.5 kV^−^; capillary temperature, 400 °C; S-lens RF level, 65; and auxiliary gas heater temperature, 350 °C. In the first analysis, aimed at comprehensive lipid profiling, samples were acquired in full scan, DDA, and DIA modes.

- For full MS scans, the resolution was set to 70,000 (at *m/z* 200), the AGC target to 1 x 10^6^, the maximum injection time to 100 ms, the scan range to *m/z* 113–1700, and data type to centroid.
- For DDA acquisition, the same MS1 settings were used, while MS/MS parameters were: resolution 70,000, AGC target 1 x 10^6^, maximum injection time 200 ms, loop count 5, isolation window 0.4 *m/z* (offset 0.4 *m/z*), collision energies 30 and 40 eV, minimum AGC target 1000, dynamic exclusion 5 s, and centroid data type. For DIA acquisition, the full scan parameters were identical to those of the MS1 acquisition, while MS/MS settings included resolution 70,000, AGC target 1 x 10^6^, maximum injection time 200 ms, collision energy 35 eV, scan range *m/z* 80– 1200, and centroid data type. An additional DIA method was performed in two scan ranges: *m/z* 50–850 and 800– 2000. In the second analysis, focused on confirming the new annotations proposed by LipidRT, iterative DDA acquisitions were carried out using the corresponding inclusion lists.

### Datasets used for validation

The NIST® Standard Reference Material® 1950 human plasma (SRM 1950) was used as the common biological reference across all validation experiments (Supplementary Table 11). Four independent RPLC-HRMS datasets acquired on different LC-MS platforms were included in this study:

- QExactive_DS1: the in-house dataset was generated using a Orbitrap Q Exactive^−^Plusmass spectrometer (Thermo Fisher Scientific) and comprised full scan, data-dependent acquisition (DDA), and data-independent acquisition (DIA) acquisitions (as described in Sample preparation and LC-MS analysis sections). This dataset was processed using different softwares as detailed in Supplementary Table 11.
- Agilent_DS2: the Agilent 6545 QTOF dataset published by Martinez *et al*. 2024^10^, which was processed using LipidMS v3.1^13^ followed by LipidRT.
- ZenoTOF_DS3: a SCIEX ZenoTOF 7600 dataset system originally published by Martinez *et al*. 2025^14^. Feature detection and alignment were performed using *xcms*^18^, followed by lipid annotation using an adaptation of the LipidMS annotation workflow compatible with *xcms* and SCIEX data. The resulting annotations were then processed with LipidRT.
- Agilent_DS4: the SRM 1950 datset using an Agilent 6546 QTOF distributed with MS-DIAL 4.0 and originally described by Tsugawa *et al*. 2020^8^, which was processed using MS-DIAL 5.5^12^ followed by LipidRT.

### Data Analysis

#### Data pre-processing and annotation

Raw LC-MS files were converted to mzXML or mzML format using msConvert (ProteoWizard v3.0.10800)^19^ for processing with LipidMS, *xcms* and LipidRT, or to ABF format using ABF Converter for processing with MS-DIAL, prior to data analysis. Depending on the dataset, lipid annotations were generated using LipidMS, MS-DIAL, Lipid Data Analyzer (LDA) or an *xcms*/LipidMS workflow before applying LipidRT (Supplementary Table 11).

For Lipid Data Analyzer (LDA, version 2.11.2)^7^, lipid annotation was performed using the glycerophospholipid, diacylglycerol and triacylglycerol databases together with the fragmentation rule sets distributed by the LDA developers. The RT post-processing option was enabled.

- For datasets processed with LipidMS v3.1.1^13^ (QExactive_DS1 and Agilent_DS2), peak picking was performed using : dmzagglom 10, drtagglom 300, drtclust 100 for MS1 and 200 for MS/MS, minpeak 5 for MS1 and 4 for MS/MS (3 for both in iterative DDA runs), drtgap 5, drtminpeak 10, drtmaxpeak 100 for MS1 and 200 for MS/MS, recurs 5 for MS1 and 10 for MS/MS, sb 3 and sn 3 for MS1, sb 2 and sn 2 for MS/MS, minint 1000 for MS1 and 100 for MS/MS, weight 2 for MS1 and 3 for MS/MS, and isotopic mass and RT tolerances (dmzIso and drtIso) 10 and 5, respectively. Batch processing (alignment and grouping) was performed using dmzalign 5, drtalign 30, span 0.4, minsamplesfracalign 0.7, dmzgroup 5, drtagglomgroup 30, drtgroup 15, and minsamplesfracgroup 0.25. Lipid annotation was then performed with precursor and product ion mass tolerances of 5 and 10 ppm, respectively, an RT tolerance of 5 s, and a coelution cutoff of 0.7.
- For datasets processed with MS-DIAL 5.5^12^ (QExactive_DS1 and Agilent_DS4), precursor and fragment mass tolerances were set to 0.005 Da and 0.01 Da, respectively. Lipid annotation required a matched spectrum score of at least 70%. Retention time was used only as a scoring criterion, and an RT tolerance of 1 min was used during peak alignment. All remaining parameters were left at their default values.
- For the ZenoTOF_DS3, feature detection and alignment were performed using *xcms*^18^. Chromatographic peaks were detected with the centWave algorithm using 25 ppm mass accuracy, peak widths of 5–50 s (MS1) and 3–50 s (MS/MS), signal-to-noise thresholds of 5 (MS1) and 2 (MS/MS), and prefilter settings of (5,100) for MS1 and (3,50) for MS/MS. Peak grouping was performed using the PeakDensity algorithm with a bandwidth of 5 s, a *minFraction* of 0.9 before retention time correction and 0.15 afterwards, and a bin size of 0.01 Da. Retention time correction was performed using the PeakGroups algorithm with LOESS smoothing (*span* = 0.4), followed by gap filling using ChromPeakArea. Lipid annotation was subsequently performed using an adaptation of the LipidMS annotation workflow with precursor and fragment mass tolerances of 10 and 15 ppm, respectively, an RT tolerance of 3 s, monoisotopic isotope confirmation enabled, isotopic peaks removed, and a minimum MS/MS intensity threshold of 100. DIA spectral reconstruction was performed using a minimum chromatographic correlation of 0.7 and a maximum RT difference of 3 s.

#### Application of LipidRT workflow to SRM 1950 datasets

Subsequently, the LipidRT workflow was applied to all the resulting SRM 1950 datasets using the default parameters (*distC_filter* = 5, *minSpecies_filter* = 5, gamma distribution and *rtscorethr* = 0.5) to remove suspicious annotations, followed by manual inspection using the integrated Shiny application. Missing annotations were then predicted using the default parameters (gamma distribution, *dmz* = 5 ppm, *rtscorethr* = 0.5, *distC_predict* = 5, *minSpecies_predict* = 5 and *rt_thr* = 30 s), and the resulting candidates were likewise reviewed manually.

#### Validation of proposed candidates by iterative DDA

Newly proposed lipid annotations were finally evaluated by combining LipidMS v3.1^13^ and MS-DIAL 5.5^12^ and manual inspection using mass and RT tolerances of 0.005 Da and 15 s, respectively.

#### Comparison of annotations across four SRM 1950 datasets

Lipid annotations were compared across the four SRM 1950 datasets analyzed in this study: the in-house acquired QExactive_DS1 and three publicly available dataset: Agilent_DS2, ZenoTOF_DS3 and Agilent_DS4 (Supplementary Table 11). Because these datasets were acquired and processed using different analytical workflows, RT values were not directly comparable and were therefore not used for matching.

For comparisons at the unique species level, annotations with different GroupIDs were considered distinct species, whereas multiple ambiguous annotations assigned to the same GroupID were counted only once. Species were matched across datasets according to lipid subclass, CN and DB. When multiple chromatographic isomers with the same sum composition were detected, the number of matches between two datasets was defined by the minimum number of isomers detected in either dataset, with any remaining isomers considered unique to the corresponding dataset.

For comparisons at the unique sum composition level, each unique combination of lipid subclass, CN and DB was counted only once, irrespective of the number of chromatographic isomers detected.

Species counts included all lipid classes analyzed in positive ionization mode (ESI(+)), except FA, LPI, PI, PG, PS and CL, for which negative ionization mode (ESI(−)) data were used.

## Data availability

The raw data required to reproduce Fig. 2b are available at Zenodo repository (https://zenodo.org) with accession number 23725495. https://doi.org/10.5281/zenodo.23725495.

The rest of the data required to reproduce the results shown in Fig. 2 or in the Supplementary Figures are provided as Supplementary Tables.

## Code availability

LipidRT R package is available via CRAN (https://CRAN.R-project.org/package=LipidRT) and the source code is available at https://github.com/maialba3/LipidRT.

## Author Contributions

M.I.A.B., J.C.G.C. and A.L. conceptualized and designed the research related to the development of LipidRT. H.C., A.G. and C.B. conceptualized and designed the research related to the elution rules governing plasmalogen chromatographic behavior. C.R.M, M.I.A.B, H.C. and M.R. performed the experiments. C.R.M and M.I.A.B. wrote the code and analyzed the data; M.I.A.B., H.C., A.G., J.C.G.C. and A.L. wrote the manuscript. C.R.M, M.I.A.B, H.C, M.R., O.J.V., J.C., A.G., C.B., J.C.G.C. and A.L. discussed the results and revised the manuscript.

