## Supplementary Text for "LipidRT: leveraging retention time coherence for robust lipid annotation in reversed-phase LC-MS-based lipidomics"

### Table of Contents


| Supplementary Notes | |
|---|---|
| **Supplementary Note 1.** Limitations of RT-based modeling when no prior filtering is applied. | iii |
| **Supplementary Note 2.** Lipid classes and fatty acyl/alkyl/alkenyl chains covered by LipidRT. | vi |
| **Supplementary Note 3.** Implementation of RT-based rules for discrimination of plasmanyl (O-) and plasmenyl (P-) glycerophospholipids (PL(O/P-)). | x |
| **Supplementary Note 4.** In silico validation of the automatic LipidRT workflow. | xv |
| **Supplementary Note 5.** Application of LipidRT workflow to lipid annotations in the SRM 1950 obtained with LipidMS. | xvii |
| **Supplementary Note 6.** Application of LipidRT workflow to lipid annotations in the SRM 1950 human plasma obtained with various LC-MS platforms and software. | xxiv |
| **Supplementary Figures** | |
| **Supplementary Fig. 1.** RT-based modeling and prediction of novel candidates using the approach described in the original LDA publication. | v |
| **Supplementary Fig. 2.** Implementation of RT-based rules for discrimination of plasmanyl (O-) and plasmenyl (P-) glycerophospholipids (PL(O/P-)). | xiii |
| **Supplementary Fig. 3.** Models for ESI(+) for the QExactive_DS1 (related to Fig. 2b). | xix-xx |
| **Supplementary Fig.4.** Models for ESI(–) for the QExactive_DS1 (related to Fig. 2b). | xxi-xxii |
| **Supplementary Fig. 5.** Details on filtered/proposed annotations for the QExactive_DS1 (related to Fig. 2b). | xxiii |
| **Supplementary Figure 6.** Lipid annotations for the 1950 SRM human plasma in various in house generated and publicly available datasets (related to Fig. 2c). | xxvi |
| **Supplementary Tables** | |
| **Supplementary Table 1.** Lipid annotations and RT-based filtered and proposed annotations related to Supplementary Fig. 1. | Excel file |
| **Supplementary Table 2.** Adducts included by default in LipidRT. | viii |
| **Supplementary Table 3.** Lipid classes and subclasses included by default in LipidRT | ix |
| **Supplementary Table 4.** Fatty acyl chains included by default LipidRT. | ix |
| **Supplementary Table 5.** Summary of solvents and concentrations of commercial and prepared stock solutions of authentic plasmalogens standards. | xiv |
| **Supplementary Table 6.** Authentic plasmalogen lipid standard composition and final concentrations of the standard (left) and spiking (right) mixtures. | xiv |
| **Supplementary Table 7.** Complete list of the initial candidates and validation results for filtering (related to Fig. 2a). | Excel file |
| **Supplementary Table 8.** Complete list of the initial candidates and validation results for prediction (related to Fig. 2a). | Excel file |
| **Supplementary Table 9.** Lipid annotations and RT-based models related to Supplementary Fig. 3 and 4. | Excel file |
| **Supplementary Table 10.** RT predicted annotations and MS/MS confirmed identities related to Fig. 2a. | Excel file |
| **Supplementary Table 11.** Description of SRM 1950 datasets used in the manuscript. | Excel file |
| **Supplementary Table 12.** Lipid annotations for all the datasets included in Fig.2c. | Excel file |
| **Supplementary Table 13.** Comparison on filtered lipid identities upon the application of the automatic LipidRT workflow alone or in combination with manual revision. | xxv |

**Supplementary Note 1. Limitations of RT-based modeling when no prior filtering is applied**.

Retention time (RT) modeling has become an increasingly important strategy for improving confidence in lipid annotation. Among the available tools, Lipid Data Analyzer (LDA) incorporates one of the most advanced RT-based post-processing workflows [7]. LDA performs targeted lipid annotation using predefined lipid databases together with lipid class-specific fragmentation rules. When RT post-processing is enabled, lipid annotations confirmed by fragmentation rules are used to fit a dataset-specific RT model. Annotations whose observed RT deviates by more than four times the root mean square error (RMSE) from the fitted regression are removed, the model is refitted, and the final regression is subsequently used to estimate the expected RT of candidate lipids that could not be confirmed by MS/MS evidence alone.

To evaluate the influence of the initial annotation quality on RT modeling, we reproduced the RT model described in the original LDA publication in-house in R using the published regression equation [$RT(x,y) = A*(1-B/x)+C*e^{(-D*y+E*x)}+F$]. The same implementation was applied to the annotation results generated by the different software packages evaluated in this study (i.e. LDA [7]., LipidMS[13] and MS-DIAL[12] ), allowing the influence of the input annotation quality on model performance to be assessed independently of the regression algorithm itself (Supplementary Figure 1, Supplementary Table 1). The exercise was executed using PC as a representative lipid class due to the high number of PC annotations that can be obtained in ESI + using the SRM 1950 as a reference material in the Orbitrap Q Exactive™ Plus mass spectrometer (Thermo Fisher Scientific) (QExactive_DS1) using the settings described in the Methods section.

As expected, the quality of the fitted RT models depended strongly on the quality of the initial annotations used for training (Supplementary Fig. 1). When the annotation workflow generated relatively few incorrect lipid identifications, as observed for LDA and LipidMS, the fitted models exhibited low RMSE values. Consequently, the initial RT filtering step was able to remove inconsistent annotations efficiently, producing refined models with narrow acceptance ranges and accurate RT predictions. In contrast, when the annotation workflow contains a larger proportion of incorrect or ambiguous annotations, the RMSE of the fitted regression model increases, resulting in considerably wider RT acceptance intervals. Because LDA defines acceptable annotations using a fixed threshold of ±4 × RMSE, the enlarged prediction window prevented many incorrect annotations from being removed during the initial filtering step. These annotations were therefore retained when the regression model was refitted, ultimately reducing the predictive accuracy of the final model and increasing the number of putatively incorrect/suspicious candidate annotations accepted during RT-based prediction. The differences in model performance are directly reflected by the width of the RT acceptance intervals shown in Supplementary Fig. 1.

These results demonstrate that the performance of regression-based RT modeling is intrinsically linked to the quality of the annotations used for model construction. LDA performs remarkably well when applied to annotations generated by its own highly restrictive rule-based workflow, where relatively few incorrect annotations are present and RT modeling effectively refines the initial results. However, when the initial annotation set contains a larger proportion of false positives, the regression model alone becomes progressively less capable of distinguishing correct from incorrect annotations. Moreover, regression-based RT models evaluate only the statistical agreement between an observed RT and the fitted mathematical relationship. They do not incorporate prior chromatographic knowledge describing the expected behavior of lipid species in reverse-phase chromatography. Consequently, candidate annotations may satisfy the regression model while violating well-established chromatographic relationships, including the expected elution order of homologous series, lipid subclasses, plasmalogens, or neighboring lipid species differing by carbon number or degree of unsaturation. A comprehensive description of the reasons to mark some of the initial or proposed annotations as putatively incorrect/suspicious based on those structure-related rules is provided in Supplementary Table 1. Another limitation of this approach is the use of a single RMSE-based acceptance criterion across the entire regression model. In practice, RT variability is rarely constant across the chromatographic space and generally increases for larger and more unsaturated lipid species. Applying the same acceptance threshold throughout the complete RT range may therefore be unnecessarily restrictive for some regions while overly permissive for others.

LipidRT was specifically designed to overcome these limitations by integrating chromatographic knowledge into both stages of the RT modeling workflow. Before model construction, a series of structural and chromatographic rules are applied to remove annotations that are inconsistent with the expected RT behavior of lipids, independently of the regression model itself. This allows robust RT models to be generated even when the original annotation set contains a substantial number of incorrect identifications. After model fitting, candidate lipid annotations are not accepted only because their predicted RT agrees with the regression model. Instead, they must also satisfy the expected chromatographic behavior of the corresponding lipid class and neighboring lipid species. In this way, RT modeling becomes one component of a broader decision framework in which prior chromatographic knowledge acts as an orthogonal structural constraint, improving both the robustness of the fitted models and the accuracy of the final lipid annotations.

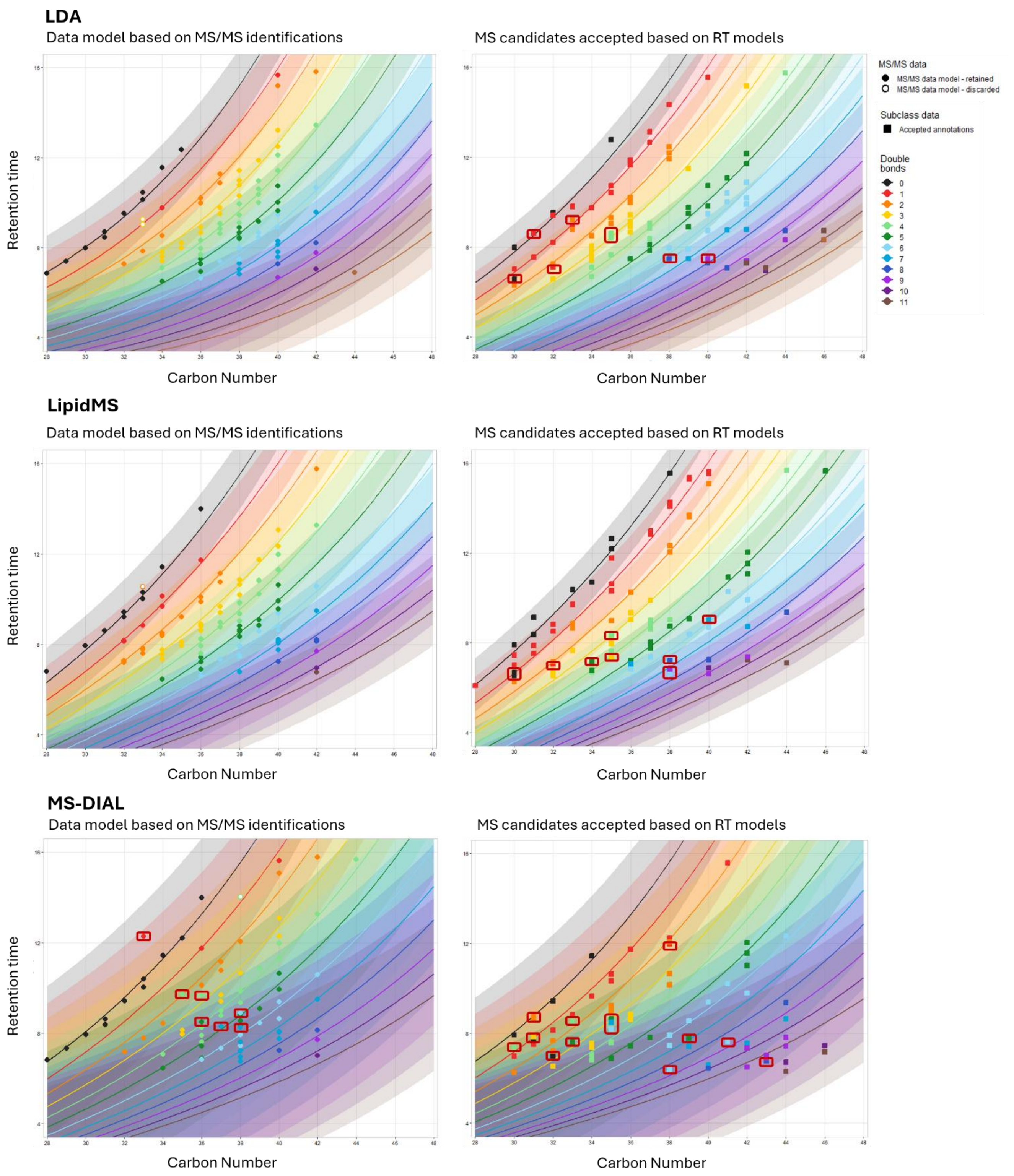


**Supplementary Fig. 1. RT-based modeling and prediction of novel candidates using the approach described in the original LDA publication** [7]. PC annotations in ESI(+) from QExactive_DS1 annotated with LDA, LipidMS or MSDIAL were modeled using the equation and acceptance criteria originally described in reference [7]. The lines describe the fitted curve and the shadow, the confidence interval. Left column: initial MS/MS based annotation, empty circles denote annotations removed based on the proposed model. Right column: proposed annotations based on the models and MS1 information. Annotations marked with a red square are suspicious annotations based on the following rule: RT[PC(CN+1:DB)]>RT[PC(CN:DB)]>RT[PC(CN:DB+1)]. Please see Supplementary Table 1 (provided as .xlsx file) for a complete list of initial and final annotations and the reasons to mark annotations as suspicious.

### Supplementary Note 2. Lipid classes and fatty acyl/alkyl/alkenyl chains covered by LipidRT.

LipidRT relies on predefined databases containing adducts, lipid classes and fatty acyl chains for candidate proposal and RT prediction. The default adducts considered for the different ionization polarities are shown in Supplementary Table 2, while the lipid classes and fatty acyl chains included by default are listed in Supplementary Tables 3 and 4, respectively. These default lists can be modified to adapt the analysis to specific experimental requirements by editing the LipidRT default database used for candidate proposal based on *m/z* values during RT prediction.

The default databases can be accessed through the Import Data tab of the Shiny application. To enable these options, users must first import a dataset using the "Choose your data import method" menu. Once data have been loaded, the customization options become available in the "Choose your lipid classes source for analysis" section. To access these options, users must select "Custom list", which enables the import and modification of the databases used for candidate proposal and RT prediction.

Importantly, only the adducts defined in the "*Adducts Table*" are considered during candidate proposal and subsequent analysis. Therefore, this table determines the set of adducts available to the software, whereas the "*File dbs Table"* specifies how those adducts are associated with individual lipid classes and fatty acyl compositions. Any adduct listed in the "*File dbs Table*" that is not present in the loaded Adducts Table will be ignored during candidate proposal and no candidates will be generated for that adduct. Consequently, adducts intended to be considered during the analysis must first be included in the "*Adducts Table*".

The Adducts Table can be accessed through the 1) "*Adducts Table*" section, where users can choose between the default configuration and a customized version. By selecting "Custom list", users can import either a CSV file or a data frame from the global environment. In addition, the "File" subsection includes a button that allows downloading the default adduct table, which can be modified and subsequently reimported to apply the desired modifications.

The custom "*Adducts Table*" must contain the columns *adducts*, corresponding to the adduct annotation, *mdiff*, indicating the mass difference associated with the adduct, *charge*, representing the ion charge state, and *n*, indicating the number of molecules involved in the adduct. All columns are mandatory and must be present using the same column names. In addition, columns must be separated by commas (","). Users can freely add or remove adducts from this table according to their analytical requirements. However, only the adducts present in the loaded "*Adducts Table*" will be considered during candidate proposal and subsequent analysis.

The "*File dbs Table"* can be customized through the 2) "*File dbs*" section by selecting "Custom list". Like the "*Adducts Table*", users may either import a new CSV file or load a data frame directly from the global environment. This allows modification of the lipid classes, adduct assignments and fatty acyl chains considered during candidate generation.

The structure of the imported table must follow the format shown in Supplementary Table 2 and contain the columns *classes*, *adducts_pos*, *adducts_neg*, *fas*, *sph*, *phyto*, *alkyl* and *vinyl*, using the same column names. The *classes* column defines the lipid classes considered during candidate proposal; new lipid classes cannot be added through this table, as LipidRT will only search for candidates belonging to the lipid classes implemented by default, although existing classes may be removed, in which case candidates from those classes will not be proposed. Furthermore, the lipid subclasses included in RT prediction can be selected directly in the *RT Prediction* tab through the *Selection of lipid subclasses for retention time prediction* option. The *adducts_pos* and *adducts_neg* columns specify the adducts associated with each lipid class in positive and negative ionization modes, respectively; although both columns are present in the default table, only the column corresponding to the polarity of the analysis is used. Therefore, users may import a table containing both polarity columns or only the column corresponding to the polarity under analysis, provided that the original column name is maintained. The *fas*, *sph*, *phyto*, *alkyl* and *vinyl* columns define the fatty acyl chain groups associated with each lipid class. In Supplementary Table 2, the cells highlighted in blue indicate the fatty acyl chain groups that constitute the lipid structure of each class and, consequently, the chain types that will be considered when proposing candidates for that class. The default fatty acyl chain compositions corresponding to these groups are listed in Supplementary Table 3. For correct import, column separators must be commas (","), whereas multiple values within a cell must be separated using the pipe character ("|"). To facilitate customization, the "File" subsection contains a download button that exports the default database (*Default Custom filedbs.csv*), which can be modified and subsequently reimported to make the changes effective.

The databases currently loaded for analysis can be inspected on the right-hand side of the Import Data tab, within the analysis information panel. In the "Lipid classes for analysis" section, separate panels are available for both the "*Adducts Table*" and the "*File dbs Table*". Each panel contains a "Show current Adducts Table" or "Show current File dbs Table" button, respectively. Clicking these buttons opens a window displaying the table currently loaded and that would be used if the analysis were executed at that moment.

**Supplementary Table 2. Adducts included by default in LipidRT.**

| Polarity | adduct | mdiff | charge | n |
|---|---|---|---|---|
| ESI(+) | M+H | 1.0078 | 1 | 1 |
| | M+H-$H_2O$ (*) | -17.0028 | 1 | 1 |
| | M+Na | 22.9892 | 1 | 1 |
| | M+$NH_4$ | 18.0344 | 1 | 1 |
| | M+H-2$H_2O$ (*) | -35.0134 | 1 | 1 |
| | 2M+$NH_4$ | 18.0344 | 1 | 2 |
| | 2M+Na | 22.9892 | 1 | 2 |
| ESI(-) | M-H | -1.0078 | -1 | 1 |
| | M-H-$H_2O$ (*) | -19.0184 | -1 | 1 |
| | M+$CH_3COO$ | 59.0139 | -1 | 1 |
| | M+HCOO | 44.9982 | -1 | 1 |
| | M+Na-2H | 20.9736 | -1 | 1 |
| | M-H-2$H_2O$ (*) | -37.029 | -1 | 1 |
| | M-$CH_3$ ($) | -15.02347 | -1 | 1 |
| | M-2H | -2.0156 | -2 | 1 |
| | 2M-H | -1.0078 | -1 | 2 |
| | M+$CH_3COOH$-$CH_3$ ($) | 44.9982 | -1 | 1 |
| | M-$H_2O$-$CH_3$ ($) | -33.03407 | -1 | 1 |
| | M-H-$CO_2$ (*) | -44.9971 | -1 | 1 |
| | M-H+$CH_3COONa$ | 80.9953 | -1 | 1 |

(*) In-source fragments resulting from neutral molecule or radical losses.

($) Putative in-source fragments; formation mechanisms may differ among lipid classes.

**Supplementary Table 3. Lipid classes and subclasses included by default in LipidRT.**

<table>
<tr><th>classes</th><th>adducts_pos</th><th>adducts_neg</th><th>fas</th><th>sph</th><th>phyto</th><th>alkyl</th><th>vinyl</th></tr>
<tr><td>Cer_d</td><td>M+H-$H_2O$|M+Na|M+H</td><td>M-H|M+$CH_3COO$|M+$CH_3COO$-$CH_3$</td><td></td><td></td><td></td><td></td><td></td></tr>
<tr><td>Cer_t</td><td>M+H-$H_2O$|M+Na|M+H</td><td>M-H|M+$CH_3COO$|M+$CH_3COO$-$CH_3$</td><td></td><td></td><td></td><td></td><td></td></tr>
<tr><td>CerP_d</td><td>M+H</td><td>M-H</td><td></td><td></td><td></td><td></td><td></td></tr>
<tr><td>CerP_t</td><td>M+H</td><td>M-H</td><td></td><td></td><td></td><td></td><td></td></tr>
<tr><td>AcylCer_d</td><td>M+H|M+H-$H_2O$|M+Na</td><td>M-H|M+$CH_3COO$</td><td></td><td></td><td></td><td></td><td></td></tr>
<tr><td>AcylCer_t</td><td>M+H|M+H-$H_2O$|M+Na</td><td>M-H|M+$CH_3COO$</td><td></td><td></td><td></td><td></td><td></td></tr>
<tr><td>HexCer_d</td><td>M+H|M+Na</td><td>M-H|M+$CH_3COO$|M+$CH_3COO$-$CH_3$</td><td></td><td></td><td></td><td></td><td></td></tr>
<tr><td>HexCer_t</td><td>M+H|M+Na</td><td>M-H|M+$CH_3COO$|M+$CH_3COO$-$CH_3$</td><td></td><td></td><td></td><td></td><td></td></tr>
<tr><td>AHexCer_d</td><td>M+H|M+Na</td><td>M-H|M+$CH_3COO$|M+$CH_3COO$-$CH_3$</td><td></td><td></td><td></td><td></td><td></td></tr>
<tr><td>AHexCer_t</td><td>M+H|M+Na</td><td>M-H|M+$CH_3COO$|M+$CH_3COO$-$CH_3$</td><td></td><td></td><td></td><td></td><td></td></tr>
<tr><td>SHexCer_d</td><td>M+H|M+Na</td><td>M-H</td><td></td><td></td><td></td><td></td><td></td></tr>
<tr><td>SHexCer_t</td><td>M+H|M+Na</td><td>M-H</td><td></td><td></td><td></td><td></td><td></td></tr>
<tr><td>SM_d</td><td>M+H|M+Na</td><td>M+$CH_3COO$|M-$CH_3$|M+$CH_3COO$-$CH_3$</td><td></td><td></td><td></td><td></td><td></td></tr>
<tr><td>SM_t</td><td>M+H|M+Na</td><td>M+$CH_3COO$|M-$CH_3$|M+$CH_3COO$-$CH_3$</td><td></td><td></td><td></td><td></td><td></td></tr>
<tr><td>FA</td><td></td><td>M-H|2M-H</td><td></td><td></td><td></td><td></td><td></td></tr>
<tr><td>HFA</td><td></td><td>M-H</td><td></td><td></td><td></td><td></td><td></td></tr>
<tr><td>Carnitine</td><td>M+H|M+Na</td><td></td><td></td><td></td><td></td><td></td><td></td></tr>
<tr><td>LPA</td><td></td><td>M-H</td><td></td><td></td><td></td><td></td><td></td></tr>
<tr><td>LPE</td><td>M+H|M+Na</td><td>M-H</td><td></td><td></td><td></td><td></td><td></td></tr>
<tr><td>LPG</td><td></td><td>M-H</td><td></td><td></td><td></td><td></td><td></td></tr>
<tr><td>LPI</td><td></td><td>M-H</td><td></td><td></td><td></td><td></td><td></td></tr>
<tr><td>LPS</td><td></td><td>M-H|M+Na-2H</td><td></td><td></td><td></td><td></td><td></td></tr>
<tr><td>LPC</td><td>M+H|M+Na</td><td>M+$CH_3COO$|M-$CH_3$|M+CH3COO-$CH_3$</td><td></td><td></td><td></td><td></td><td></td></tr>
<tr><td>MG</td><td>M+H-$H_2O$|M+ $NH_4$|M+Na</td><td></td><td></td><td></td><td></td><td></td><td></td></tr>
<tr><td>CE</td><td>2M+$NH_4$|2M+Na|M+ $NH_4$|M+Na</td><td></td><td></td><td></td><td></td><td></td><td></td></tr>
<tr><td>Sph_d</td><td>M+H</td><td>M-H</td><td></td><td></td><td></td><td></td><td></td></tr>
<tr><td>Sph_t</td><td>M+H</td><td>M-H</td><td></td><td></td><td></td><td></td><td></td></tr>
<tr><td>SphP_d</td><td>M+H</td><td>M-H</td><td></td><td></td><td></td><td></td><td></td></tr>
<tr><td>SphP_t</td><td>M+H</td><td>M-H</td><td></td><td></td><td></td><td></td><td></td></tr>
<tr><td>NAGly</td><td>M+H|M+Na</td><td>M-H</td><td></td><td></td><td></td><td></td><td></td></tr>
<tr><td>NAGlySer</td><td>M+H|M+Na</td><td>M-H</td><td></td><td></td><td></td><td></td><td></td></tr>
<tr><td>NAOrn</td><td>M+H|M+Na</td><td>M-H</td><td></td><td></td><td></td><td></td><td></td></tr>
<tr><td>NAE</td><td>M+H|M+Na</td><td>M-H</td><td></td><td></td><td></td><td></td><td></td></tr>
<tr><td>DG</td><td>M+H- $H_2O$ |M+NH4|M+Na</td><td></td><td></td><td></td><td></td><td></td><td></td></tr>
<tr><td>PC</td><td>M+H|M+Na</td><td>M+$CH_3COO$|M-$CH_3$| M+$CH_3COO$-$CH_3$</td><td></td><td></td><td></td><td></td><td></td></tr>
<tr><td>PE</td><td>M+H|M+Na</td><td>M-H</td><td></td><td></td><td></td><td></td><td></td></tr>
<tr><td>PG</td><td>M+H|M+$NH_4$|M+Na</td><td>M-H</td><td></td><td></td><td></td><td></td><td></td></tr>
<tr><td>PS</td><td></td><td>M-H|M+Na-2H</td><td></td><td></td><td></td><td></td><td></td></tr>
<tr><td>PI</td><td>M+H|M+$NH_4$|M+Na</td><td>M-H</td><td></td><td></td><td></td><td></td><td></td></tr>
<tr><td>PA</td><td></td><td>M-H</td><td></td><td></td><td></td><td></td><td></td></tr>
<tr><td>FAHFA</td><td></td><td>M-H</td><td></td><td></td><td></td><td></td><td></td></tr>
<tr><td>MGDG</td><td>M+$NH_4$|M+Na</td><td>M-H</td><td></td><td></td><td></td><td></td><td></td></tr>
<tr><td>DGDG</td><td>M+$NH_4$|M+Na</td><td>M-H</td><td></td><td></td><td></td><td></td><td></td></tr>
<tr><td>BMP</td><td>M+$NH_4$|M+Na</td><td>M-H</td><td></td><td></td><td></td><td></td><td></td></tr>
<tr><td>LNAPE</td><td>M+$NH_4$|M+Na</td><td>M-H</td><td></td><td></td><td></td><td></td><td></td></tr>
<tr><td>TG</td><td>M+$NH_4$|M+Na</td><td></td><td></td><td></td><td></td><td></td><td></td></tr>
<tr><td>CL</td><td></td><td>M-H|M+Na-2H</td><td></td><td></td><td></td><td></td><td></td></tr>
<tr><td>PE-O</td><td>M+H|M+Na</td><td>M-H|M+Na$CH_3COO$</td><td></td><td></td><td></td><td></td><td></td></tr>
<tr><td>PC-O</td><td>M+H|M+Na</td><td>M+$CH_3COO$|M-$CH_3$|M+$CH_3COO$-$CH_3$</td><td></td><td></td><td></td><td></td><td></td></tr>
<tr><td>LPE-O</td><td>M+H|M+Na</td><td>M-H|M+Na $CH_3COO$</td><td></td><td></td><td></td><td></td><td></td></tr>
<tr><td>LPC-O</td><td>M+H|M+Na</td><td>M+$CH_3COO$|M-$CH_3$|M+$CH_3COO$-$CH_3$</td><td></td><td></td><td></td><td></td><td></td></tr>
<tr><td>LPA-O</td><td></td><td>M-H</td><td></td><td></td><td></td><td></td><td></td></tr>
<tr><td>PE-P</td><td>M+H|M+Na</td><td>M-H|M+Na$CH_3COO$</td><td></td><td></td><td></td><td></td><td></td></tr>
<tr><td>PC-P</td><td>M+H|M+Na</td><td>M+$CH_3COO$|M-$CH_3$|M+$CH_3COO$-$CH_3$</td><td></td><td></td><td></td><td></td><td></td></tr>
<tr><td>LPE-P</td><td>M+H|M+Na</td><td>M-H|M+Na$CH_3COO$</td><td></td><td></td><td></td><td></td><td></td></tr>
<tr><td>LPC-P</td><td>M+H|M+Na</td><td>M+$CH_3COO$|M-$CH_3$| M+$CH_3COO$-$CH_3$</td><td></td><td></td><td></td><td></td><td></td></tr>
</table>

**Supplementary Table 4. Fatty acyl chains included by default LipidRT.**

<table>
<tr><th>fas</th><th>alkyl</th><th>vinyl</th><th>sph</th><th>phyto</th></tr>
<tr><td colspan="3">8:0|10:0|12:0|14:0|14:1|15:0|16:0|16:1|17:0|18:0|18:1|18:2|18:3|18:4|20:0|20:1|20:2|20:3|20:4|20:5|<br>22:0|22:1|22:2|22:3|22:4|22:5|22:6|24:0|24:1|24:2|24:3|24:4|24:5|24:6|26:0|28:0|30:0|32:0|34:0|36:0</td><td>16:0|16:1|18:0|<br>18:1|18:2|20:0|20:1</td><td>18:0</td></tr>
</table>

### Supplementary Note 3. Implementation of RT-based rules for discrimination of plasmanyl (O-) and plasmenyl (P-) glycerophospholipids (PL(O/P-)).

The previously established LC-MS-based lipidomic database of NIST® SRM® 1950 human plasma (SRM 1950) [10,14] contains high-confidence lipid annotations supported by collision-induced dissociation (CID) and electron impact excitation of ions from organics (EIEIO) MS/MS spectra. This database provides a reference for subsequent LC-MS lipidomic experiments, enabling lipid annotation based on accurate mass and retention time (RT) without requiring additional MS/MS acquisition. However, MS/MS-based annotation of the SRM 1950 database initially assigned PL(O/P-) pairs as alternative candidates, as these isomers cannot generally be distinguished by their MS/MS spectra.

To establish chromatographic rules for discriminating plasmanyl and plasmenyl glycerophospholipids, fifteen commercially available plasmenyl (plasmalogens) PE(P), LPE(P), PC(P), and LPC(P) authentic chemical lipid standards were analyzed as pure standards and after spiking into pooled human plasma using the same RP-UHPLC-ESI(+)-QTOF-MS workflow employed for construction of the SRM 1950 database[10]. More details are provided in the Material and Methods. Matching the plasmalogens standards with their corresponding endogenous features enabled experimental assignment of the plasmenyl species, thereby resolving the ambiguity in the MS/MS annotations. The complementary chromatographic feature was consequently assigned as the corresponding plasmanyl isomer (Supplementary Fig. 2a).

Comparison of the experimentally confirmed PL(O/P-) pairs with the corresponding endogenous diacyl analogues demonstrated that the *sn*-1 double bond position and the functional group (carbonyl *vs* ether) produce a reproducible RT shift in reversed-phase liquid chromatography (Supplementary Fig. 2a,b). These observations were generalized into the following chromatographic rule:

$$RT[PL(O\text{-}CN{:}DB+1)] < RT[PL(CN{:}DB)] < RT[PL(P\text{-}CN{:}DB)] < RT[PL(O\text{-}CN{:}DB)]$$

which was incorporated into the LipidRT algorithm to support annotation filtering and discrimination of previously unresolved PL(O/P-) species in complex biological samples (Supplementary Fig. 2c). This chromatographic rule expands and generalizes previous observations regarding the chromatographic behavior of plasmanyl and plasmenyl glycerophospholipids in RPLC[20].

Application of the algorithm to the complete SRM 1950 Agilent_DS2 first identified eight MS/MS-derived ether glycerophospholipid annotations that were inconsistent with the expected chromatographic behavior and were therefore excluded prior to RT modeling. The validated annotations were subsequently used to generate RT models, enabling discrimination of 50 plasmanyl/plasmenyl species based on their RT behavior, comprising 34 PC and 16 PE lipids (Supplementary Fig. 2d). Together, the chromatographic rules and RT modeling provided a robust and scalable strategy for PL(O/P-) classification in complex biological samples, and the only available alternative when MS/MS validation of the proposed candidates is not possible.

**Materials and methods related to Supplementary Note 3**

Analytical Standards and Reagents

Ultrapure Milli-Q $H_2O$ was obtained using a Milli-Qplus185 system (18 MΩ·cm, Millipore, Bedford, MA, USA). Methanol (MeOH), acetonitrile (ACN), ethanol (EtOH) and chloroform ($CHCl_3$) and isopropanol (IPA), all LC-MS grade, were obtained from CARLO ERBA Reagents (Sabadell, Spain). HPLC-grade methyl tert-butyl ether (MTBE) (99.99%, CHROMASOLV® Plus), ammonium fluoride ($NH_4F$) (ACS reagent, ≥ 98%), and glacial acetic acid (ACS reagent, ≥ 99.8%) were purchased from Sigma-Aldrich (Steinheim, Germany). Analytical grade ammonia solution (28%, GPR RECTAPUR®) was obtained from VWR Chemicals (Radnor, PA, USA). SPLASH® LIPIDOMIX® Mass Spec Standard, a stable mixture of 14 deuterium-labeled lipids, and 10 authentic chemical lipid standards –LPC(P-16:0/0:0), LPC(P-18:0/0:0), LPE(P-18:0/0:0), PC(P-16:0/18:1), PC(P-18:0/18:1), PC(P-18:0/20:4), PC(P-18:0/22:6), PE(P-18:0/18:1), PE(P-18:0/20:4), PE(P-18:0/22:6)– were purchased from Avanti Polar Lipids Inc. (Alabaster, AL, United States). The other 5 authentic lipid standards, including PC(P-16:0/16:0), PC(P-16:0/18:2), PC(P-16:0/20:4), PC(P-18:0/22:4), PE(P-18:0/22:4), were acquired from Cayman Chemical (Ann Arbor, MI, USA).

Lipid standard mixture preparation

Stock solutions of solid authentic plasmenyl (plasmalogens) standards were prepared by dissolution in either IPA or IPA/$H_2O$ mixtures according to compound solubility. The solvent composition and final concentrations of the prepared stock solutions, together with the commercially supplied standard solutions and their corresponding solvents, are summarized in Supplementary Table 5.

Subsequently, five plasmalogens standard mixtures, each containing three authentic standards from the LPC, LPE, PC, and PE subclasses, were prepared at a final concentration of 5 ppm by dilution of either prepared stock solutions or commercially supplied standard solutions using IPA as diluent.

Based on preliminary signal responses obtained from the analysis of plasmalogen standard mixtures and a pooled plasma, an additional set of five spiking standard mixtures was prepared. For plasmalogens detected endogenously, spiking concentrations were adjusted to achieve approximately a two-fold increase relative to endogenous abundance after spiking. To minimize analytical interferences, all plasmalogens standard mixtures were designed to avoid isomeric and isobaric overlaps between compounds within the same mixture. The composition and final concentrations of the standard and spiking mixtures are detailed in Supplementary Table 6.

Sample preparation: Lipid Extraction and Spiking

Lipids from a pooled plasma sample were extracted following an all-in-one single-phase lipid extraction method based on a monophasic solvent mixture composed of MeOH/MTBE (1:1, v/v) to cover the isolation of polar and non-polar lipids. Plasma samples were thawed on ice and homogenized by vortexing for 2 min.

For pooled plasma, 50 µL of plasma was mixed with 175 µL of MeOH, followed by the addition of 175 µL of MTBE. For spiked plasma, 50 µL of plasma was mixed with 165 µL of MeOH, followed by the addition of 175 µL of MTBE and 10 μL of the corresponding spiking standard mixture. MeOH volume was adjusted to 165 μL to maintain a constant final solvent volume across all samples.

SPLASH® LIPIDOMIX® internal standard was thawed on ice, vortexed to ensure homogeneity, and 10 μL was added before extraction. The resulting mixture was vortexed for 30 min at room temperature, followed by sample centrifugation at 16,100 x g for 15 min at 15 °C. Finally, 200 μL of the resulting supernatant was transferred to LC Chromacol (Agilent Chromacol) vials with insert and centrifuged at 2,000 x *g* for 5 min at 15 °C before analysis. Blank solutions were prepared containing only Milli-Q $H_2O$ and the solvent mixture, and the same procedure was followed.

Analytical Conditions

Pooled plasma, spiked plasma lipid extracts, and the authentic endogenous lipid standard mixtures were analyzed using an Agilent 1290 Infinity II UHPLC system coupled to an Agilent Revident QTOF Mass Spectrometer (Agilent Technologies, Santa Clara, CA, USA) equipped with a Dual Agilent Jet Stream Electrospray source (Dual AJS ESI).

For the RP-UHPLC analysis, an optimized method already reported in previous lipidomic studies was implemented and employed[10].

Pooled plasma, spiked plasma lipid extracts, and the authentic endogenous lipid standard mixtures were injected in positive and negative ionization mode analysis with volumes of 0.5 μL and 1 μL, respectively, using an Agilent 1290 Infinity II Multisampler system kept at 15 °C to maintain compound stability and prevent lipid precipitation. An Agilent InfinityLab Poroshell 120 EC-C18 (100 mm × 3 mm × 2.7 μm) reversed-phase column and an Agilent InfinityLab Poroshell 120 EC-C18 (3 mm × 5 mm × 2.7 μm) guard column were used for lipid separation, and the temperature was maintained at 50 °C. Mobile phases used for both ionization modes were (A) 10 mM $CH_3COONH_4$ and 0.2 mM $NH_4F$ in $H_2O$/MeOH (9:1, v/v) and (B) 10 mM $CH_3COONH_4$ and 0.2 mM $NH_4F$ in ACN/MeOH/IPA (2:3:5, v/v/v). The flow rate was set at 0.6 mL/min. The gradient during the chromatographic analysis began with 70% of B at 0 – 1 min, 86% B at 3.5 – 10 min, and 100% B at 11 – 17.00 min. The initial conditions were restored by min 17.10 followed by a 1.90 min re-equilibration, leading to a total running time of 19 min. The strategy used for multiwash involved a mixture of MeOH/IPA (1:1, v/v) and a wash time of 15 s, followed by a mixture of A/B (30:70, v/v) to assist in reaching the initial conditions.

Acquisition was performed in full MS scan in positive ionization mode (ESI(+)). The Agilent Revident QTOF Mass Spectrometer equipped with a Dual AJS ESI ion source was set with the following parameters: 175 V fragmentor, 65 V skimmer, 3500 V capillary voltage, 1000 V nozzle voltage, 750 V octopole radio frequency voltage, 11 L/min drying gas flow, 290 °C gas temperature, 40 psi nebulizer gas pressure, 11 L/min sheath gas flow, and 370 °C sheath gas temperature.

A solution consisting of two reference mass compounds was infused into the system throughout the whole analysis at a flow rate of 40 μL/min and a reference nebulizer pressure of 8 psi for mass correction. The reference masses used were *m/z* 121.0509 (purine detected as $[C_5H_4N_4+H]^+$) and *m/z* 922.0098 (HP-0921 detected as

[$C_{18}H_{18}O_6N_3P_3F_{24}+H]^+$). Data acquisition was done using Agilent MassHunter LC/MS Acquisition Software v 12.1.98.0 (Agilent Technologies, Waldbronn, Germany), firmware v 1.2.38. The MS operated in full scan mode (MS1 level) and was scanned from *m/z* 100 – 1700 at a scan rate of 1 spectra/s.

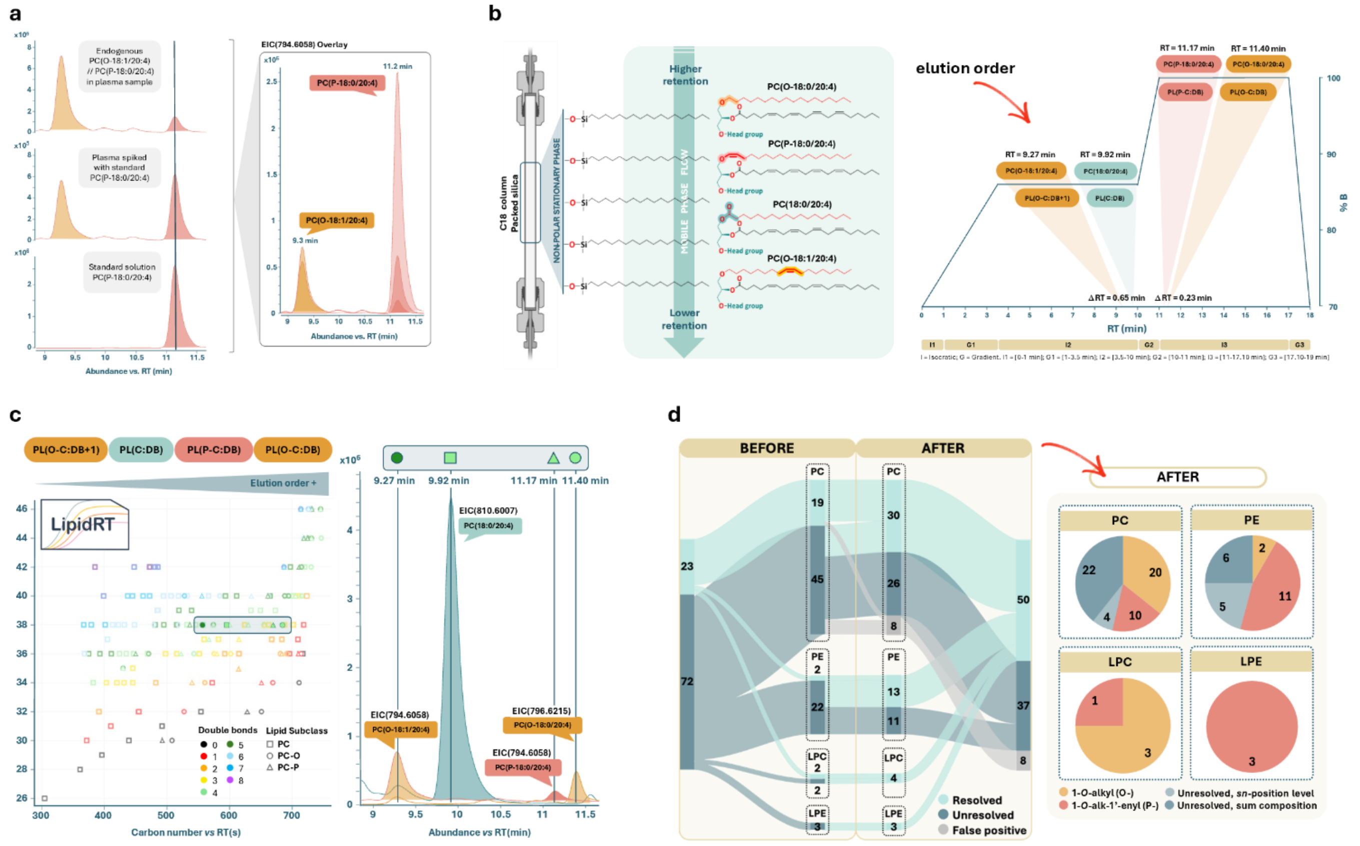


**Supplementary Fig. 2. Implementation of RT-based rules for discrimination of plasmanyl (O-) and plasmenyl (P-) glycerophospholipids (PL(O/P-)). a**, Experimental confirmation of PL(O/P-) pairs using the previously established SRM 1950 Agilent_DS2 dataset. Representative extracted ion chromatograms (EIC) of endogenous PC(O-18:1/20:4) and PC(P-18:0/20:4) in pooled human plasma, spiked plasma, and pure standard solution illustrate the assignment of previously unresolved PL(O/P-) isomers. **b**, Chromatographic basis of PL(O/P-) discrimination. The functional group and position of the *sn*-1 double bond determine the retention order in RPLC, with plasmenyl species eluting after their plasmanyl counterparts of identical molecular composition. Relative retention behavior is illustrated using representative PC species. **c**, Generalization of the experimentally observed chromatographic behavior into a rule-based algorithm implemented in LipidRT. The algorithm propagates experimentally established retention behavior to support discrimination of unresolved PL(O/P-) annotations in complex biological sample. **d**, Application of the algorithm to the SRM 1950 lipidomic database from Agilent_DS2 (published table). RT-based filtering removed chromatographically inconsistent annotations prior to RT modeling, after which the resulting RT models enabled discrimination of previously unresolved plasmanyl and plasmenyl glycerophospholipids across the PC and PE subclasses.

**Supplementary Table 5. Summary of solvents and concentrations of commercial and prepared stock solutions of authentic plasmalogens standards.**

| Lipid name | Solvent | Concentration (ppm) |
|---|---|---|
| **Prepared stock solutions** | | |
| PC(P-16:0/18:1) | IPA | 1000,00 |
| PC(P-16:0/16:0) | IPA | 1000,00 |
| LPC(P-16:0/0:0) | IPA/$H_2O$ (20:1, v/v) | 952,38 |
| LPC(P-18:0/0:0) | IPA/$H_2O$ (20:1, v/v) | 952,38 |
| LPE(P-18:0/0:0) | IPA/$H_2O$ (21:4, v/v) | 800,00 |
| **Commercial standard solutions** | | |
| PC(P-18:0/18:1) | $CHCl_3$ | 1000,00 |
| PC(P-18:0/20:4) | $CHCl_3$ | 1000,00 |
| PC(P-18:0/22:6) | $CHCl_3$ | 1000,00 |
| PE(P-18:0/18:1) | $CHCl_3$ | 1000,00 |
| PE(P-18:0/20:4) | $CHCl_3$ | 1000,00 |
| PE(P-18:0/22:6) | $CHCl_3$ | 1000,00 |
| PC(P-16:0/20:4) | EtOH | 1000,00 |
| PC(P-18:0/22:4) | EtOH | 5000,00 |
| PE(P-18:0/22:4) | EtOH | 1000,00 |
| PC(P-16:0/18:2) | EtOH | 5000,00 |

**Supplementary Table 6. Authentic plasmalogen lipid standard composition and final concentrations of the standard (left) and spiking (right) mixtures.**

| Standard mixtures | | Spiking standard mixtures | |
|---|---|---|---|
| **Common name** | **Final concentration (ppm)** | **Common name** | **Final concentration (ppm)** |
| **Mix 1** | | **Mix 1** | |
| PE(P-18:0/18:1) | 5 | PE(P-18:0/18:1) | 2 |
| PC(P-18:0/22:6) | 5 | PC(P-18:0/22:4) | 2 |
| PC(P-18:0/18:1) | 5 | LPC(P-18:0/0:0) | 2 |
| **Mix 2** | | **Mix 2** | |
| PC(P-18:0/20:4) | 5 | PC(P-18:0/22:6) | 200 |
| PC(P-16:0/18:2) | 5 | PC(P-16:0/20:4) | 200 |
| PE(P-18:0/20:4) | 5 | PE(P-18:0/22:4) | 200 |
| **Mix 3** | | **Mix 3** | |
| PC(P-16:0/20:4) | 5 | PC(P-18:0/18:1) | 3,5 |
| PC(P-16:0/18:1) | 5 | LPE(P-18:0/0:0) | 3,5 |
| PE(P-18:0/22:4) | 5 | LPC(P-16:0/0:0) | 3,5 |
| **Mix 4** | | **Mix 4** | |
| LPE(P-18:0/0:0) | 5 | PC(P-18:0/20:4) | 20 |
| PE(P-18:0/22:6) | 5 | PE(P-18:0/22:6) | 20 |
| LPC(P-16:0/0:0) | 5 | PC(P-16:0/16:0) | 20 |
| **Mix 5** | | **Mix 5** | |
| PC(P-16:0/16:0) | 5 | PC(P-16:0/18:2) | 90 |
| PC(P-18:0/22:4) | 5 | PE(P-18:0/20:4) | 72 |
| LPC(P-18:0/0:0) | 5 | PC(P-16:0/18:1) | 54 |

### Supplementary Note 4. In silico validation of the automatic LipidRT workflow.

To evaluate the filtering and prediction steps implemented in LipidRT, we used the SRM 1950 database reported by Martínez *et al.* 2024[10] (Agilent_DS2). Only data acquired in positive ionization mode were considered for this validation as it provides the widest coverage of the lipidome in the reference database. Given that the reported database contains several adducts for each lipid and that experimentally the most common situation is that more abundant lipids can be detected as various co-eluting adducts while lower abundant ones can be detected as the single most abundant adduct, we decided to simulate adduct distribution so that only the 2 most commonly found adducts for each lipid class were retained and in each iteration lipids were randomly assigned to be represented by 1 or 2 co-eluting adducts following the proportion 50:50, based on the expected distribution of previously analysed lipidomics datasets for the SRM 1950.

Validation of RT-based filtering

To evaluate whether LipidRT correctly removes chromatographically inconsistent annotations, artificial annotations were introduced into the curated reference dataset. In each experiment, 5%, 15%, 25%, 35% or 45% of the original annotations were randomly selected and duplicated. For each duplicated annotation, the expected RT was defined as the mean RT of the reference annotations belonging to the same lipid subclass and having the same sum composition. The RT of the duplicated annotation was then artificially displaced from this expected value by adding or subtracting, with equal probability, the RT variability (defined as the standard deviation) observed for that lipid subclass multiplied by a predefined factor (1.5, 2, 2.5, 3, 3.5 or 4). Thus, increasing displacement factors generated artificial annotations with progressively greater deviations from the expected chromatographic behavior while preserving the original lipid identity. Each combination of artificial annotation proportion and RT displacement factor was independently repeated ten times, with annotations randomly selected and the direction of the RT displacement randomly assigned in each iteration.

The resulting datasets were processed with the automated LipidRT filtering workflow, and filtering performance was quantified as the proportion of artificial annotations removed. LipidRT increasingly rejected annotations as their chromatographic inconsistency increased. Across artificial annotation proportions, approximately 70% of artificial annotations were removed at an RT displacement of 1.5-fold the subclass-specific variability, increasing progressively to approximately 87% at a 4-fold displacement, being ≈81% the mean value across RT displacements and proportion of incorrect features introduced. In contrast, the proportion of artificial annotations introduced had a comparatively limited effect on filtering performance (Fig. 2a). These results show that LipidRT preferentially removes annotations according to the magnitude of their inconsistency with the expected chromatographic behavior rather than simply responding to the overall proportion of potentially incorrect annotations in the dataset. Regarding the particular annotations that were not removed during the filtering process, classes with the lowest number of annotated species were more prone to keeping incorrect annotations than classes with the highest number of annotations (supplementary table 7). In general, those lipid classes with less than 20 lipids had errors above the mean value, while species with 20 or more lipids had errors below the mean value. It is worth mentioning that lipid classes with restrictions in the fatty acyl chain composition of the chains (e.g. Cer or SM) were more prone to errors (Supplementary Table 7) as the number of comparisons between particular lipid species to evaluate whether an annotation should be maintained or removed is lower and thus less restrictive (please see Methods and Fig. 1b for details about the filtering algorithm implemented within LipidRT).

Validation of lipid prediction

Prediction performance was evaluated independently by removing the annotations from 5%, 10%, 15%, 20%, 25%, 30%, 35% or 40% of the high-confidence dataset (evenly distributed across lipid classes) while retaining the corresponding features. The complete automatic LipidRT workflow was then applied, and the identities predicted for these features were compared with their original curated annotations. This procedure was repeated ten times for each removal proportion, with annotations randomly selected in each iteration. Predictions were considered correct when the predicted and original annotations matched in lipid subclass and sum composition of the top 1 predicted annotation or any other top n annotation with an RTscore > 0.5 that differs less than 0.1 from the top 1 annotation.

LipidRT recovered approximately 89% of the removed annotations across all proportions evaluated, with little variation in recovery rate as the proportion of removed annotations increased from 5% to 40%. Importantly, among features for which LipidRT proposed an annotation, approximately 98% were correctly assigned. The difference between recovery and prediction accuracy reflects the conservative behavior of the workflow: when the remaining annotations did not provide sufficient chromatographic support for a candidate, LipidRT preferentially left the feature unannotated rather than assigning a poorly supported lipid identity. Thus, this controlled validation showed that LipidRT can recover a substantial proportion of missing annotations while maintaining high prediction accuracy even when the proportion of available reference annotations is progressively reduced. In general, the mistakes were evenly distributed across lipid classes, for RT-based modeling and prediction related classes are combined when the number and distribution of the lipid species do not allow to build a model for each lipid class/subclass. However, there was a clear pattern to not predict species that corresponded to the lowest or highest limit of the fitted distributions as those are the regions with the highest discrepancy between the observed and the predicted RT values (Supplementary Table 8).

### Supplementary Note 5. Application of LipidRT workflow to lipid annotations in the SRM 1950 obtained with LipidMS.

To evaluate the complete LipidRT workflow under controlled conditions, the in-house QExactive_DS1 was processed using LipidMS followed by LipidRT. Since LipidMS applies stringent fragmentation rules for lipid annotation, this dataset provides a high-confidence starting point for assessing the impact of RT-based filtering and prediction independently of major annotation errors.

<u>RT-based filtering</u>

Application of the LipidRT filtering workflow substantially improved the consistency of the initial annotation set by removing chromatographically implausible lipid annotations and resolving ambiguities between isobaric/isomeric species (Fig. 2b, Supplementary Fig. 3-5, Supplementary Table 9). Most discarded annotations corresponded to isobaric/isomeric lipid species whose MS/MS spectra alone were insufficient to confidently distinguish between alternative structures, but whose retention times violated established chromatographic relationships. Consequently, the filtering procedure reduced ambiguities between structurally related lipids (Fig.2b, Supplementary Fig. 5). The filtering procedure reduced < 15 % the number of unique species while reducing the number of ambiguous identifications by ≈ 80%.

Although the workflow is fully automated, manual inspection through the LipidRT Shiny application remains advisable, particularly for lipid subclasses represented by only a small number of annotated species. In these cases, the lack of neighboring lipids differing by one carbon atom or one double bond limits the amount of chromatographic information available to evaluate RT consistency. While the implemented filtering rules remain effective under these conditions, occasional incorrect annotations may persist or isolated correct annotations may be discarded because insufficient neighboring species are available to establish the expected chromatographic behavior. Such cases are readily identified during visual inspection and can be corrected before RT model construction.

Overall, RT-based filtering generated a high-confidence annotation set with substantially fewer ambiguous and incorrect annotations, providing the high-confidence annotation set required for reliable dataset-specific RT modeling and prediction.

<u>RT-based prediction</u>

The curated annotation set was subsequently used to construct dataset-specific RT models and predict lipid species from unidentified MS1 features. Application of the prediction workflow produced a substantial increase in lipidome coverage while maintaining high annotation confidence (Fig. 2a, Supplementary Fig. 3-5).

Overall, ≈20% of the proposed annotations corresponded to additional adducts of previously identified lipids, while the remaining annotations correspond to novel isomeric species of previously annotated lipids or previously unreported sum CN:DB compositions, thus demonstrating that LipidRT expands lipidome coverage beyond increasing annotation redundancy (Fig. 2a and Supplementary Fig. 5).

Considering unique lipid species defined by the combination of sum composition and retention time (with RT tolerance of 5 s), the number of annotated lipids increased from 409 to 854 (> 100% increase) following application of the complete LipidRT workflow. When chromatographic isomers were grouped and annotations were collapsed according to sum composition only, the number of unique lipid compositions increased from 318 to 573 (≈ 80% increase) (Fig. 2a and Supplementary Fig. 5).

Inspection of the newly predicted annotations revealed that they predominantly filled discontinuities within homologous CN and DB series, producing chromatographically coherent lipid distributions (Supplementary Fig. 3-4). This observation suggests that many of these annotations correspond to low-abundance lipid species that were already detected at the MS1 level but lacked sufficiently informative MS/MS spectra for confident identification during the initial annotation process. Rather than predicting isolated lipid species, LipidRT preferentially recovered structurally consistent members of existing lipid series, thereby improving the completeness of the reconstructed lipidome.

Experimental validation of the predicted lipids

To experimentally evaluate the proposed annotations, iterative DDA analyses were performed using inclusion lists generated from the LipidRT predictions, taking into consideration exclusively novel unique species, but not new adducts of previously annotated lipids, and annotated using LipidMS and MSDIAL followed by manual inspection using mass and RT tolerances of 0.005 Da and 15 s, respectively. 238 MS/MS spectra acquired using DDA corresponded to features annotated by LipidRT. In 193 cases (133 in ESI(+) and 60 in ESI(–)) the MS/MS information confirmed the top1 annotation proposed by LipidRT. In 4 cases (1 in ESI(+) and 3 in ESI(–)) the MS/MS spectra matched the top2 annotation proposed by LipidRT and 32 (13 in ESI(+) and 19 in ESI(–)) correspond to unresolved plasmalogens, where the MS/MS spectra can confirm the annotation, but the combination of MS/MS spectra and RT is insufficient to distinguish between PL-O and PL-P. Finally, only in 9 cases (5 in ESI(+) and 4 in ESI(–)) the obtained spectra validated a lipid annotation different that the one proposed by LipidRT. Thus, the proportion of incorrect annotations was < 4% (Fig. 2a, Supplementary Table 10).

These results demonstrate that combining knowledge-based chromatographic filtering with dataset-specific RT modeling enables robust prediction of previously unidentified lipid species while maintaining high annotation accuracy. Importantly, the final validation step implemented in LipidRT, in which predicted annotations are evaluated not only according to their agreement with the regression model but also against independent chromatographic constraints, proved essential for preventing biologically implausible predictions and maintaining high confidence in the final annotation set.

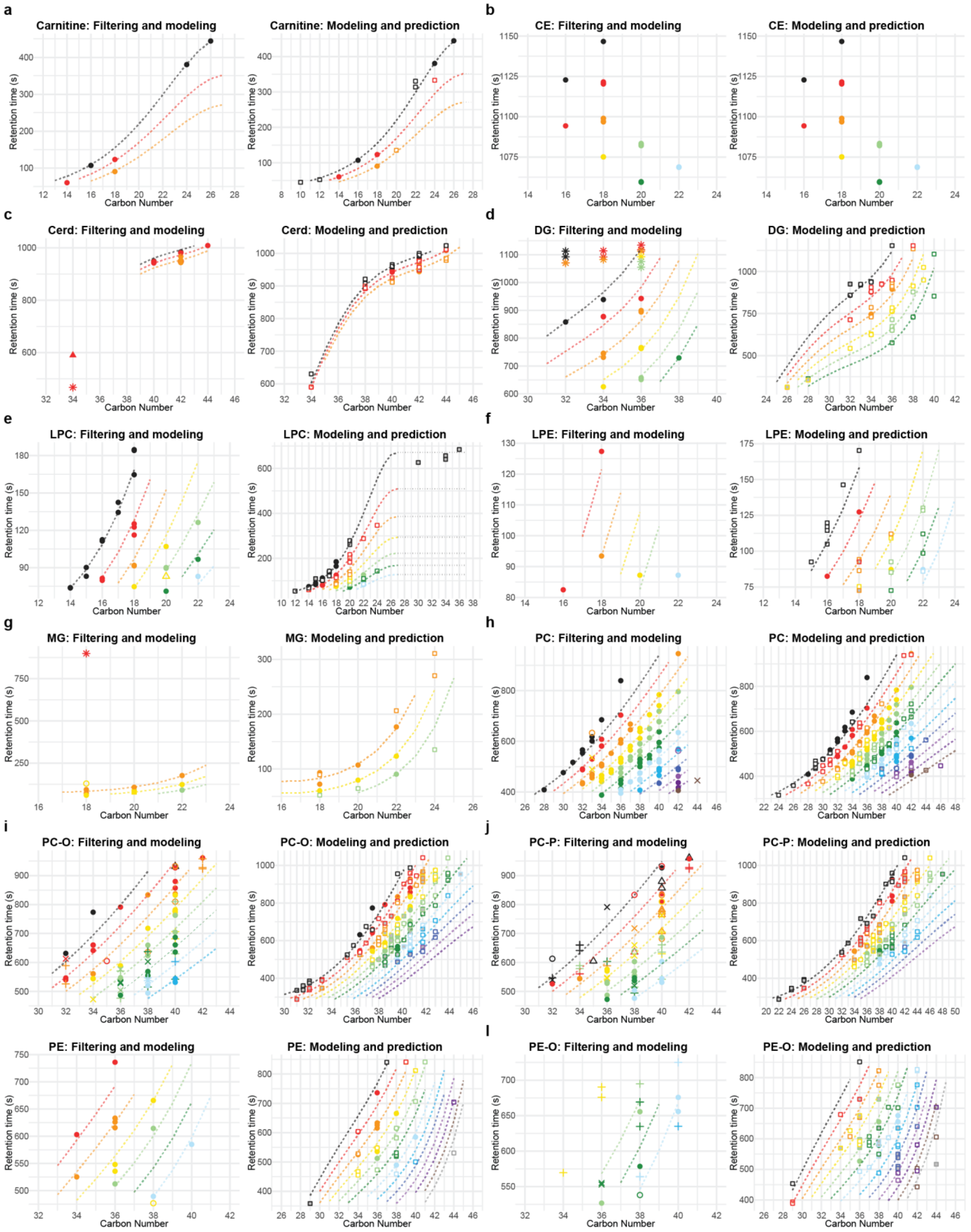


**Supplementary Fig. 3. Models for ESI(+) for the QExactive_DS1 (related to Fig. 2b).**

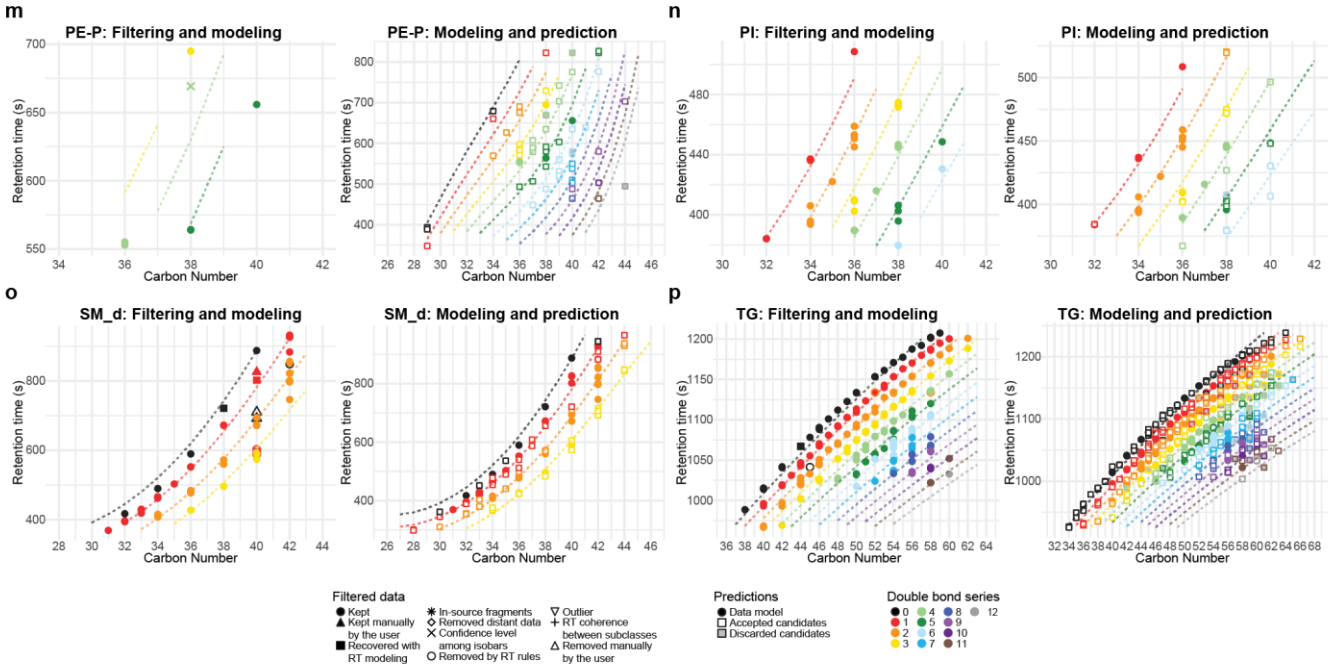


**Supplementary Fig. 3. Models for ESI(+) for the QExactive_DS1 (related to Fig. 2b)(continued).**

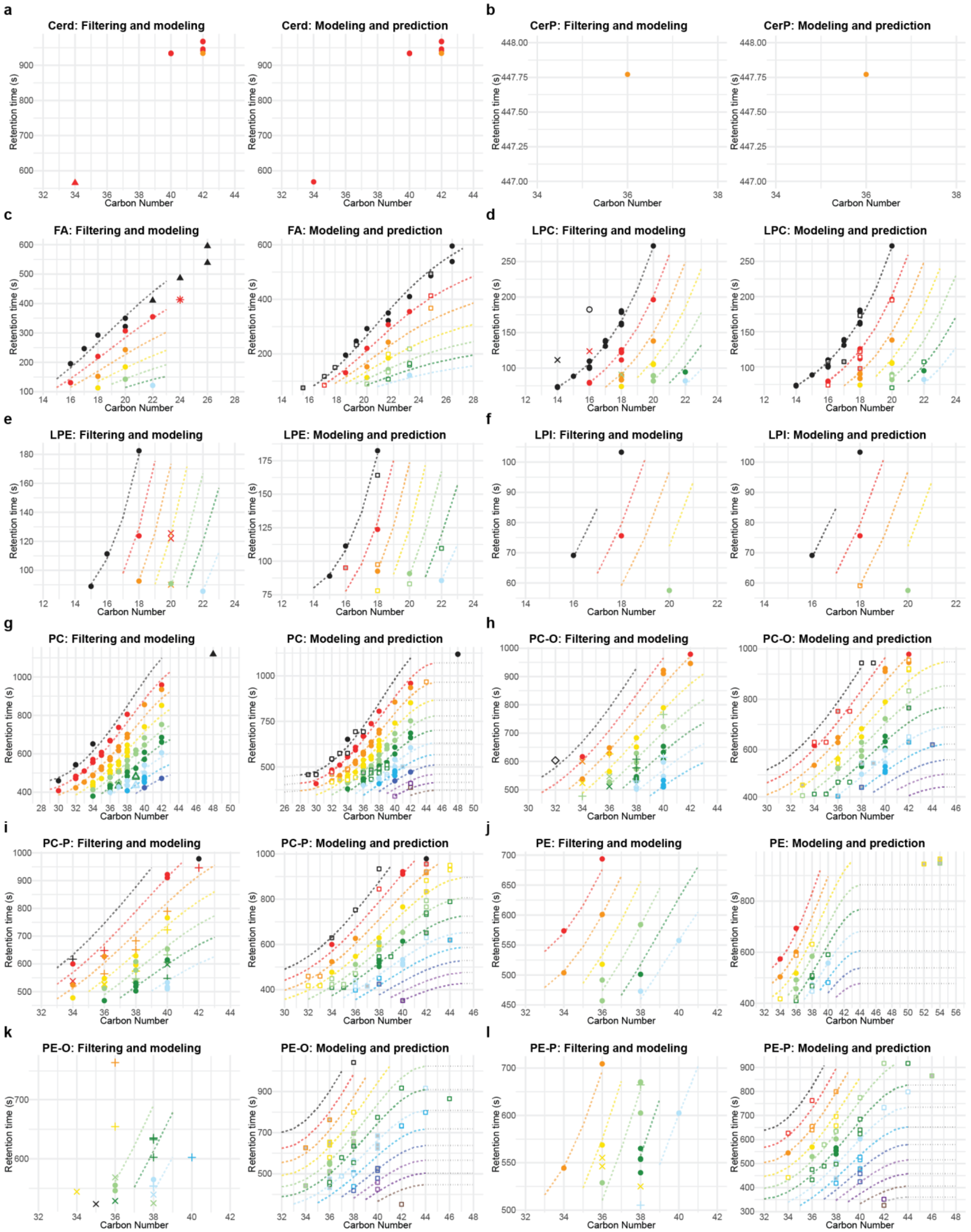


**Supplementary Fig.4. Models for ESI(–) for the QExactive_DS1 (related to Fig. 2b).**

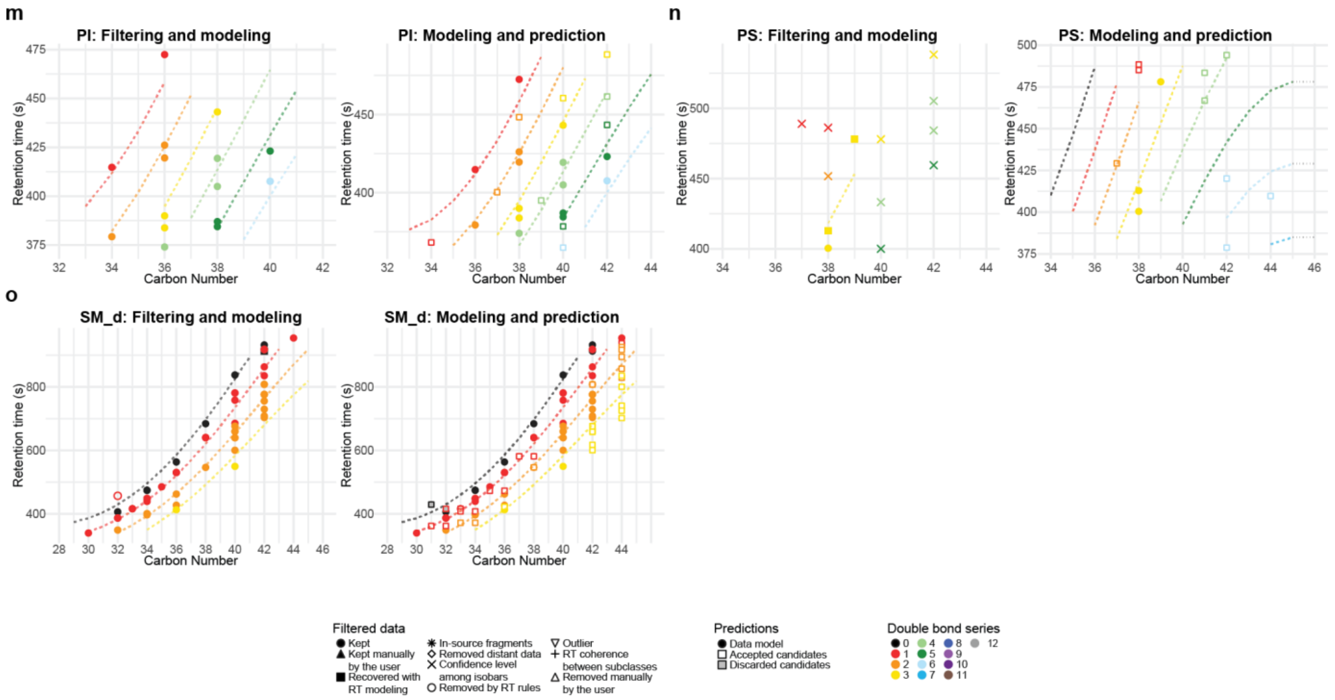


**Supplementary Fig.4. Models for ESI(–) for the QExactive_DS1 (related to Fig. 2b) (continued).**

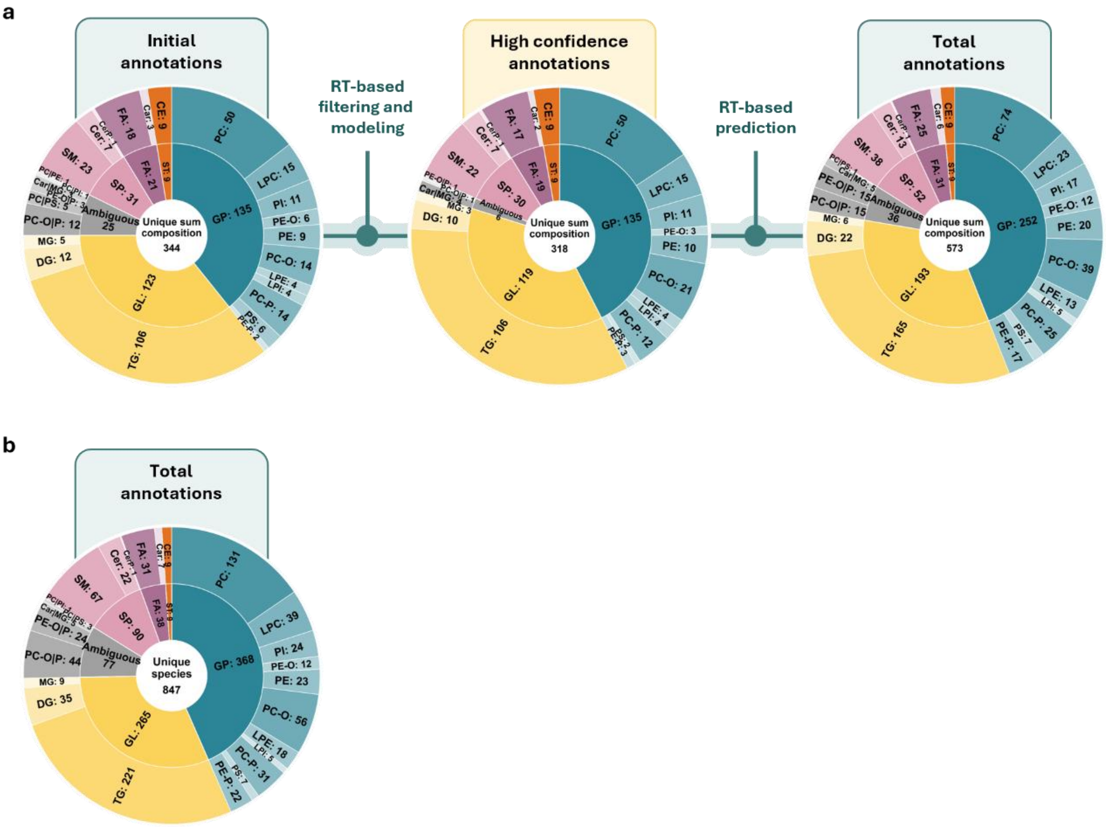


**Supplementary Fig. 5. Details on filtered/proposed annotations for the QExactive_DS1 (related to Fig. 2b). a**, Initial, high confidence and total (after RT-based filtering, modeling and prediction) number of identified lipids based on unique sum composition (unique CN:DB combinations, without taking into consideration isomers). **b**, Total number of annotated lipids (unique species, taking into consideration isomers) after RT-based filtering, modeling and prediction.

### Supplementary Note 6. Application of LipidRT workflow to lipid annotations in the SRM 1950 human plasma obtained with various LC-MS platforms and software.

We applied LipidRT to an in-house generated (QExactive_DS1) and various published [8,10,14] (Agilent_DS2, ZenoTOF_DS3 and Agilent_DS4) lipidomics datasets from SRM 1950 human plasma acquired on multiple RPLC-HRMS platforms (Thermo Scientific Q Exactive™ Plus, Agilent 6545/6546 QTOF and SCIEX ZenoTOF 7600 system) (Supplementary Table 11). In all the cases the annotations obtained with both positive and negative ionization modes were combined to obtain a unified list of annotated, unique and unique sum composition lipid species to facilitate interplatform comparison given that each dataset used a different chromatographic separation and even different gradients and/or chromatographic conditions were used for positive and negative ionization modes for each platform. The complete initial, filtered and predicted annotations for the four datasets can be found at Supplementary Table 12.

First, we decided to use the four datasets, covering multiple experimental and data processing options, to compare the results of the automatic LipidRT workflow vs the automatic in combination with manual curation LipidRT workflow (Supplementary Table 13). The automatic filtering reduced the number of lipid ids by approximately 18% (18 ± 5%, min. 9%, max. 27%). This number is consistent with the estimation of incorrect annotations initially described for LipidMS 3.0[13]. Further inspection of the data taking advantage of LipidRT ShinyApp, where the models can be easily visualized and manually curated, lead to an additional reduction of approximately 6% of the retained annotations (6 ± 6%, min. 2%, max 18%). As expected based on previous experience, in datasets annotated with LipidMS 3.0[13] manual curation only further reduced approx. 3% of the retained annotations whereas the dataset annotated with MS-DIAL 5.5[12], which applies less stringent criteria, manual revision reduced approx. 16% of the initial annotations. LipidRT does not aim to replace manual curation but to provide an assisted and intuitive framework that facilitates the refinement of lipid annotations by combining the automatic application of a series of well-stablished structure-related RT-based constrains with a graphic interface that enables expert-user curation in a more efficient way.

After the application of filtering step of LipidRT, the ZenoTOF_DS3 had the highest number of unique species and unique sum composition species across the four datasets included in the study (Fig.2c and Supplementary Fig.6), which demonstrates the speed and sensitivity of this platform compared to the other ones. The same chromatographic method was used for the Agilent_DS2 and the ZenoTOF_DS3 datasets, the first providing the lowest number of initial annotations. The Agilent_DS4 dataset was obtained using a very similar method (using a 19-min separation instead of a 17-min one), and the QExactive_DS1 used the longest chromatographic method, with a 24-min separation. Interestingly, while the ZenoTOF_DS3 dataset contained the largest number of species not shared with the other datasets (Fig.2c and Supplementary Fig.6) before RT-prediction (183 of 568 species, ≈32%), as expected based on its large initial coverage of the lipidome, the Agilent_DS4 had the highest proportion of lipids not shared with the rest of the datasets (155 of 390 species, ≈40%). This can be due to the use of MS-DIAL 5.5 software for the annotation of this dataset compared to the use of LipidMS for the rest of the datasets, as MS-DIAL 5.5 covers a larger number of lipid classes and fatty acyl chains than LipidMS and uses a different strategy for lipid annotation.

Upon the application of RT-based lipid prediction there was an increase of approx. ≈80-100% in unique species and approx. ≈65-80% in unique sum composition for all the datasets except for the ZenoTOF_DS3 where the increase was ≈30% for both parameters as the initial lipidome coverage, with high quality MS/MS data, was large enough to prevent a massive detection of lipid species with good MS1 detection, but low quality MS/MS spectra.

The application of RT-based prediction resulted in ≈60% increase in unique species and ≈40% in unique sum composition. While the increase in lipid annotations common to one, two or three datasets increased in the range of ≈30 - 65% for unique species and ≈10 - 30% for unique sum compositions, the increase in lipids common to the four datasets increase ≈140-150% for both unique species and unique sum composition, thus highlighting the capacity of LipidRT to homogenize lipid coverage across lipidomic datasets originated in highly divergent experimental and computational conditions.

**Supplementary Table 13. Comparison on filtered lipid identities upon the application of the automatic LipidRT workflow alone or in combination with manual revision.**

<table>
<tr><th colspan="3"></th><th>QExactive_DS1</th><th>Agilent_DS2</th><th>ZenoTOF_DS3</th><th>Agilent_DS4</th></tr>
<tr><td rowspan="3">ESI(+)</td><td colspan="2">Initial Annotations</td><td>717</td><td>519</td><td>1148</td><td>732</td></tr>
<tr><td rowspan="2">Removed annotations</td><td>Automatic workflow</td><td>135</td><td>84</td><td>313</td><td>136</td></tr>
<tr><td>Manual revision</td><td>16</td><td>13</td><td>35</td><td>83</td></tr>
<tr><td rowspan="3">ESI(–)</td><td colspan="2">Initial Annotations</td><td>435</td><td>111</td><td>274</td><td>354</td></tr>
<tr><td rowspan="2">Removed annotations</td><td>Automatic workflow</td><td>94</td><td>13</td><td>26</td><td>74</td></tr>
<tr><td>Manual revision</td><td>14</td><td>2</td><td>9</td><td>52</td></tr>
</table>

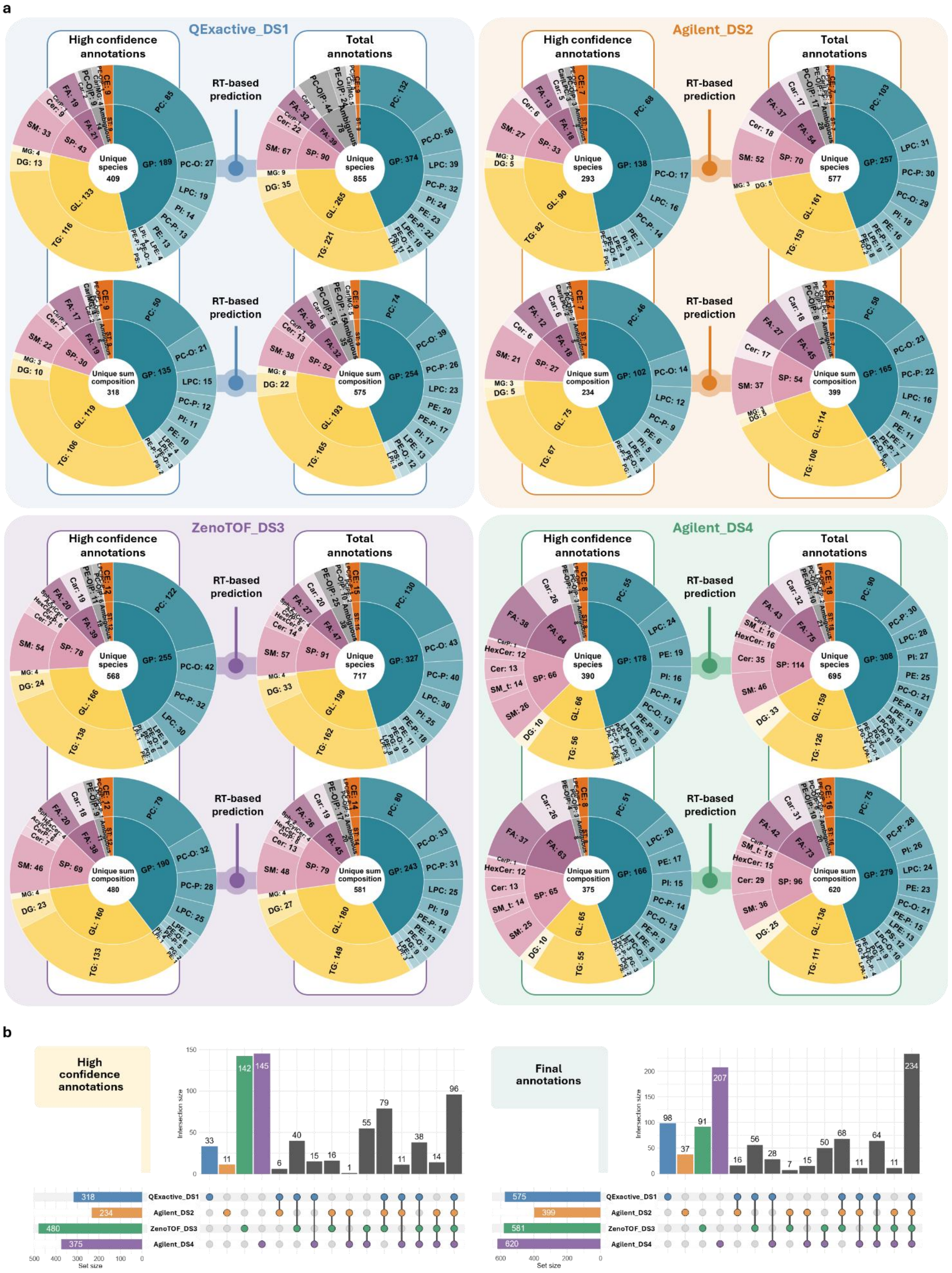


**Supplementary Figure 6. Lipid annotations for the 1950 SRM human plasma in various in house generated and publicly available datasets (related to Fig. 2c). a**, Number of annotations for each of the datasets included in the comparison after the filtering (high confidence annotations) and the prediction steps (total annotations). **b**, Comparison of the lipidome coverage across the four LC-MS datasets in terms of unique sum composition species.
